# Genomic epidemiology of the cholera outbreak in Yemen reveals the spread of a multi-drug resistance plasmid between diverse lineages of *Vibrio cholerae*

**DOI:** 10.1101/2022.08.24.504966

**Authors:** Florent Lassalle, Salah Al-Shalali, Mukhtar Al-Hakimi, Elisabeth Njamkepo, Ismail Mahat Bashir, Matthew J. Dorman, Jean Rauzier, Grace A. Blackwell, Alyce Taylor-Brown, Mathew A. Beale, Ali Abdullah Al-Somainy, Anas Al-Mahbashi, Khaled Almoayed, Mohammed Aldawla, Abdulelah Al-Harazi, Marie-Laure Quilici, François-Xavier Weill, Ghulam Dhabaan, Nicholas R. Thomson

## Abstract

The humanitarian crisis in Yemen led in 2016 to the biggest cholera outbreak documented in modern history, with more than 2.5 million suspected cases to date. In late 2018, epidemiological surveillance showed that *V. cholerae* isolated from cholera patients had turned multi-drug resistant (MDR). We generated genomes from 260 isolates sampled in Yemen between 2018 and 2019 to identify a possible shift in circulating genotypes. 84% of *V. cholerae* isolates were serogroup O1 belonging to the seventh pandemic El Tor (7PET) lineage, sublineage T13 – same as in 2016 and 2017 – while the remaining 16% of strains were non-toxigenic and belonged to divergent *V. cholerae* lineages, likely reflecting sporadic gut colonisation by endemic strains. Phylogenomic analysis reveals a succession of T13 clones, with 2019 dominated by a clone that carried an IncC-type plasmid harbouring an MDR pseudo-compound transposon (PCT). Identical copies of these mobile elements were found independently in several unrelated lineages, suggesting exchange and recombination between endemic and epidemic strains. Treatment of severe cholera patients with macrolides in Yemen from 2016 to early 2019 coincides with the emergence of the plasmid-carrying T13 clone. The unprecedented success of this genotype where an SXT-family integrative and conjugative element (SXT/ICE) and an IncC plasmid coinhabit show the stability of this MDR plasmid in the 7PET background, which may durably reduce options for epidemic cholera case management. We advocate a heightened genomic epidemiology surveillance of cholera to help control the spread of this highly-transmissible, MDR clone.

## Introduction

Since 2016, Yemen has seen the largest epidemic of cholera ever recorded. This occurred against the backdrop of a civil war turned international conflict and famine which together fueled extensive population movement, with more than 4 million people internally displaced by the end of 2020^1^. The Electronic Disease Early Warning System (eDEWS), a surveillance programme coordinated by the Ministry of Public Health and Population of Yemen (MPHP) in Sana’a tasked with monitoring the epidemic^2^, had recorded a total of almost 2.4 million suspected cholera cases up until August 2019^3^. These cases exhibited a seasonal profile, with peaks in July 2017 and September 2018 (16,000 and 50,000 cases per week, respectively)^3^. The lower reported case incidence in 2018 was ascribed to the mass vaccination campaign led by the World Health Organization (WHO) and United Nation Children’s Fund (UNICEF), who delivered the oral cholera vaccine (OCV) to 540,000 people in August 2018 (387,000 at follow-up in September) in targeted districts in Aden, Hudaydah and Ibb governorates^4,5^. Notwithstanding this focussed vaccination campaign, cholera cases were recorded nationwide in 2019, peaking at over 30,000 cases per week. Despite the mass vaccination campaign, case numbers declined at a slower rate than in previous years^3^.

Pandemic cholera is caused by discrete phylogenetic lineages of the bacterium *Vibrio cholerae* that are associated with epidemic spread, and carry lipopolysaccharide O-antigens of serogroups O1 or O139. The large majority of epidemic strains associated with cholera outbreaks from the last 60 years belong to the seventh pandemic El Tor (7PET) lineage of *V. cholerae* O1, which swept the planet in three pandemic waves^6^. We previously used genomic epidemiology to show that the first two waves of the cholera outbreak in Yemen (2016 and 2017) were driven by a single clonal expansion^7^ belonging to Wave 3 of the global 7PET lineage and had an Ogawa serotype. This indicated the Yemen outbreak was seeded by a single international transmission event linked to the 7PET sublineage involved in the thirteenth recorded intercontinental introduction of cholera (T13)^7^.

Our ongoing surveillance activities in Yemen found that the fluctuating peaks in incidence in Yemen were accompanied by a sudden change in the antibiotic susceptibility profile reported by the reference laboratory at the MPHP in Sana’a. Whilst strains isolated in 2016-2018 were sensitive to most of the antibiotics usually used for the treatment of cholera (excepting quinolones, where reduced suceptibility to ciprofloxacin prevented the use of this antibiotic as a single dose treatment), by 2019, resistance was observed for multiple drugs including third generation cephalosporins, macrolides (including azithromycin) and cotrimoxazole. Whilst the main treatment for cholera is rehydration therapy, antibiotics can be used to limit the volume and duration of the acute watery diarrhoea, and reduce the risk of transmission^8–10^. In Yemen, macrolides were used extensively up to early 2019 to treat moderate to severe cases of cholera in pregnant woman and children, the latter forming the large majority of cases^11^. Multiple drug resistance (MDR) in *V. cholerae* is strongly associated with the acquisition of mobile genetic elements (MGEs) such as SXT-family integrative and conjugative elements (SXT/ICE) or plasmids of the incompatibility type C (IncC, formerly known as IncA/C2; ref. 12), which often carry and disseminate antimicrobial resistance (AMR) gene cargo^13^.

We hypothesised that the MDR phenotype seen in the Yemen *V. cholerae* isolates from 2019 could be explained either by gain of resistance (either through *de novo* mutations or acquisition of resistance-conferring MGEs) in the previously susceptible 7PET-T13 *V. cholerae* strain already circulating in Yemen, or through the replacement of that strain with locally or globally derived MDR strain(s). Distinguishing between these hypotheses is important for understanding the ongoing dynamics of cholera in Yemen, and will be important for cholera control strategies. We therefore applied genomic epidemiology approaches to determine the molecular basis for the observed switch to the MDR phenotype and its link to global and local evolutionary dynamics of pandemic cholera. In doing so, we highlight the role of globally circulating MGEs in making an epidemic pathogen resistant to multiple drugs and subsequently reducing treatment options. We also show that these MGEs and their cargo AMR genes were repeatedly exchanged among diverse *V. cholerae* lineages found in Yemen.

## Results

### Sampling of *V. cholerae* in Yemen in 2018 and 2019

The National Centre of Public Health Laboratories (NCPHL) in Sana’a, the capital city, received 6,311 and 3,225 clinical samples collected from suspected cholera patients, in 2018 and 2019 respectively. Of these, 2,204 (35%) and 2,171 (67%) were confirmed to be positive for *V. cholerae* O1 by culture (identification based on biochemical tests and detection of Ogawa and Inaba serotypes; Table S7; Figure S1). Among the 1,642 *V. cholerae* isolated at the NCPHL from January to October 2018, 623 were tested for susceptibility to a range of antibiotics by the disk diffusion method, of which 620 (99.6%) were phenotypically resistant to nalidixic acid and nitrofurantoin, but otherwise sensitive to all other antimicrobials tested (Figure S2; Tables S7). In contrast, all tested *V. cholerae* isolates (*n* = 2,172) from January 2019 onwards were resistant to nalidixic acid, azithromycin, co-trimoxazole and cefotaxime (Figure S2; Tables S7), a pattern maintained up to late 2021 (WHO EMRO, personal communication). The transition in phenotype occurred during November 2018, when 159/175 (90.8%) tested isolates already showed the MDR profile. 250 of the 2018-2019 clinical *V. cholerae* isolates were randomly chosen for further characterization (Table S1). These samples originated from eight of the 21 Yemen governorates, comprising 71 out of 333 districts (Table S1), with 101 samples collected in 2018 (from mid-July to late October) and 149 in 2019 (from late February to late April and from early August to mid-October). In addition, ten environmentally-derived strains were isolated from sewerage in Sana’a in October 2019 (Table S1).

Extended antibiotic sensitivity testing of these 260 isolates at NCPHL and Institut Pasteur (IP) (Figure S3) reflected the phenotypic switch to MDR observed in the wider sample set, further showing that all tested 2019 strains were resistant to ampicillin, cefotaxime, nalidixic acid, azithromycin, erythromycin and co-trimoxazole (Tables S1, S2; Supplementary text).

### Phylogenetic diversity of the *V. cholerae* isolated in Yemen in 2018 and 2019

We isolated a single colony for 240 out of the 260 *V. cholerae* isolates indicated above, and multiple independent colony picks for the remaining 20 isolates, for a total of 281 isolates on which we performed whole genome sequencing (Figure S3; Tables S1, S2). After quality filtering, this yielded 232 high-quality isolate genome assemblies (selecting a single isolate from each initial sample), which we combined with 650 previously published *V. cholerae* O1 and non-O1 genomes for context, totaling 882 assembled genomes (Table S4; Figure S3). We inferred a core-genome phylogeny for this genome set, which described the sequenced diversity of the *V. cholerae* species, rooted by the genomes that belong to its newly described sister species *V. paracholerae*^14^. We subdivided *V. cholerae* genomes according to their distribution in eleven crown clades of the core-gene phylogeny clades, referred to henceforth as *Vc*A to *Vc*K (Figures 1, S3; Table S4). *Vc*H contained all 7PET epidemic lineage genomes utilised in this dataset, including 663 contextual genomes, the the majority (216/232) of the Yemen 2018-2019 genomes, and all 42 previously reported 2016-2017 Yemeni genomes^7^ (Figure S4).

**Figure 1:**
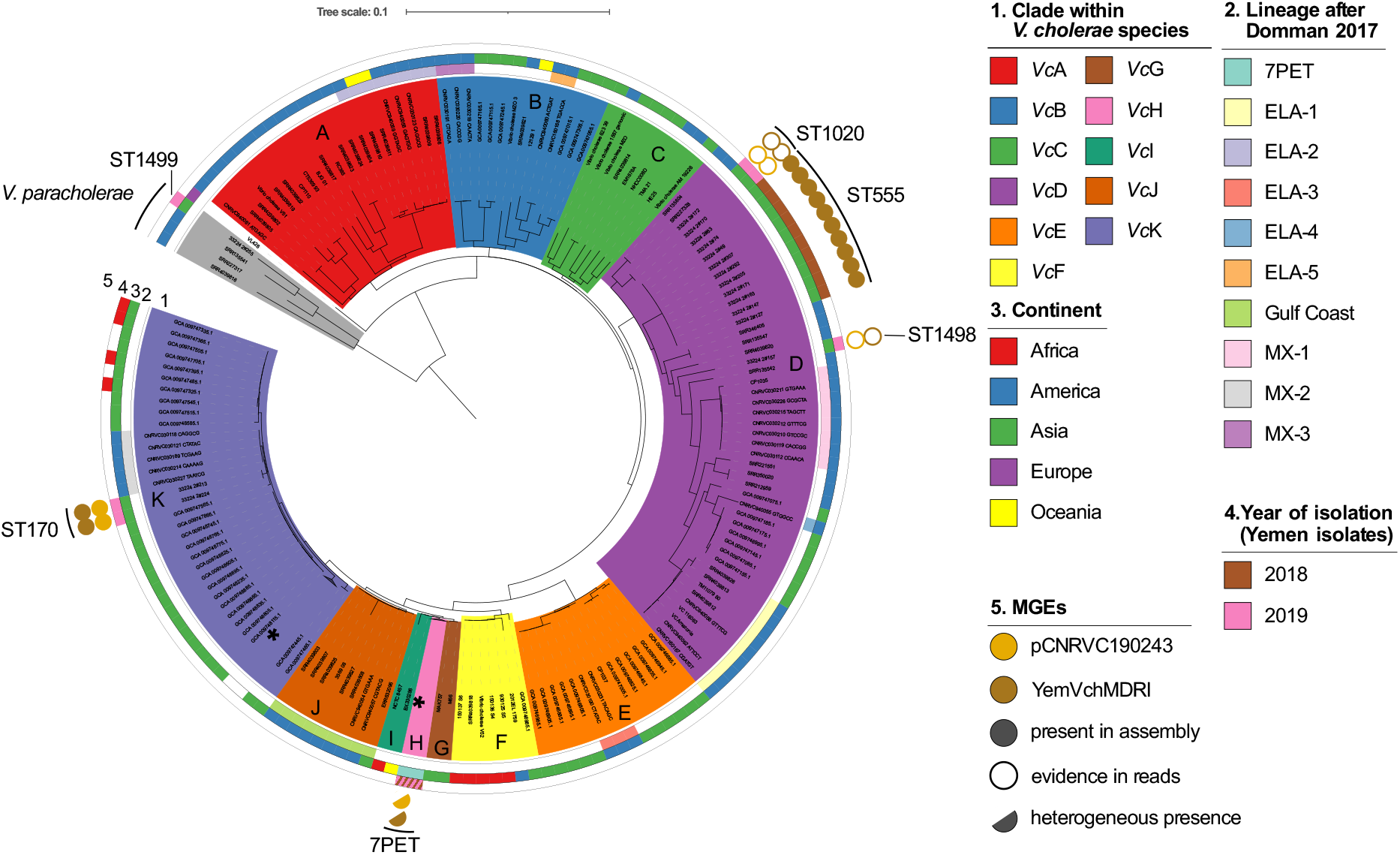
Phylogenetic diversity of *Vibrio cholerae* isolates from Yemen. Maximum-likelihood phylogeny of 882 assembled *V. cholerae* genomes based on the 37,170 SNP sites from the concatenated alignments of 291 core genes. Low-diversity clades (*Vc*H and part of *Vc*K) are collapsed and marked by black stars. Clades are highlighted with background colours (legend key 1). Coloured rings outside the tree depict the match with previously described lineages (ring 2), the geographical origin of isolates at the level of continents (ring 3), and their year of isolation when from Yemen (ring 4). Presence of parts of the plasmid pCNRVC190243 are indicated by coloured circles (ring 5 in A): IncC plasmid backbone (light brown) and the MDR pseudo-compound transposon Yem*Vch*MDRI (dark brown); full circles indicate over 70% coverage in assemblies of the reference length, hollow circles indicate 30-70% coverage in assemblies and confirmed presence based on mapped reads, with even coverage over the MGE reference sequence, while half-circles represent heterogeneous presence in a collapsed clade. Tree plots were generated with iTOL v4^16^ and adapted with Inkscape. The scale bar represents the number of nucleotide substitutions per variable site.

Whilst Yemeni *Vc*H isolates show limited genomic diversity (99.98-100.00% ANI similarity; 0 to 97 SNPs), the remaining 16 Yemeni genomes belonged to clades *V. paracholerae* (*Vpc*), *Vc*D and *Vc*K and were overall more diverse than *Vc*H isolate genomes (96.24-99.99% ANI similarity; Figure 1; Table 1); these represent “non-7PET” lineages. Based on core genome phylogeny and MLST, we found five distinct clusters within three non-7PET clades: *Vpc* (*n* = 1; novel ST1499), *Vc*D (*n* = 21; ST555, ST1020 and novel ST1498; Table S6) and *Vc*K (*n* = 2; ST170) (Figure 1).

**Table 1.** Number of *V. cholerae* isolate genomes from Yemen by year and phylogenetic lineage.

| Year | Total | Clades |  |  |  | Clusters |  |  |  |  |
| --- | --- | --- | --- | --- | --- | --- | --- | --- | --- | --- |
|  |  | *Vpc*<sup>1</sup> | *VcD*<sup>2</sup> | *VcK*<sup>1</sup> | *VcH/H.9*<sup>3</sup> |  |  |  |  | n.d.<sup>4</sup> |
|  |  |  |  |  |  | H.9.e | H.9.f | H.9.g | H.9.h |  |
| 2016 | 8 |  |  |  | 8 | 7 | 1 |  |  |  |
| 2017 | 34 |  |  |  | 34 | 29 | 5 |  |  |  |
| 2018 | 112 |  | 17 |  | 87 | 3 |  | 6 | 78 | 8 |
| 2019 | 169 | 1 | 4 | 2 | 151 |  |  | 150 | 1 | 11 |
| Total | 323 | 1 | 21 | 2 | 280 | 39 | 6 | 156 | 79 | 19 |

<sup>1</sup> Assigned based on the “882 assembled *V. cholerae* genomes” dataset
<sup>2</sup> Assigned based on the “33 mapped *VcD* genomes” dataset
<sup>3</sup> Assigned based on the 456 “mapped 7PET genomes”
<sup>4</sup> Poor quality genome data or no coverage of the bacterial genomes (e.g. in case of complete contamination by ICP1 virus genome).

Although highly clonal, phylogenetic structure within *Vc*H allowed it to be further subdivided into subclades *Vc*H.1 to *Vc*H.10 (Figure S5). All the Yemen 2016-2019 isolates fell within *Vc*H.9, which corresponds to the T13 sublineage of 7PET Wave 3 (ref. 7). We selected one representative isolate (CNRVC190243) of *Vc*H.9, and used PacBio sequencing to generate long reads in addition to the Illumina short reads obtained for all samples, which enabled us to generate a closed hybrid assembly. We subsequently used Oxford Nanopore sequencing to do the same for a *Vc*D representative isolate (CNRVC190247). To obtain greater phylogenetic resolution within *Vc*H.9, we then mapped sequencing reads to our new *Vc*H.9 CNRVC190243 reference genome to build a “mapped genome tree”. Here, together with our novel *Vc*H.9 genomes (*n* = 238), we included 218 previously published genomes that reside in this subclade and close outgroups, for a total of 456 genomes (Table S5). This approach allowed us to further subdivide *Vc*H.9 into phylogenetic clusters named *Vc*H.9.a to *Vc*H.9.h (Figure 2A). Yemeni genomes form a monophyletic group (clusters *Vc*H.9.e to *Vc*H.9.h), emerging from the genetic diversity of East African genomes (clusters *Vc*H.9.c and *Vc*H.9.d), which in turn branch out of a cluster of South Asian genomes (*Vc*H.9.b), consistent with previous observations on the origins of 7PET-T13, introduced from South Asia into Africa ^7,15^. Clusters *Vc*H.9.g and *Vc*H.9.h together comprise the majority of 2018-2019 Yemen isolates (235/281) and form a well-supported clade (94% bootstrap) that branches from within *Vc*H.9.f (Table 1). Cluster *Vc*H.9.h includes the majority of the Yemeni 7PET-T13 isolates (78/87) from 2018, with just one isolate from March 2019. In contrast, Cluster *Vc*H.9.g comprises mostly 2019 isolates (150/156), and a minority from 2018 (6/156) (Table 1). All of the 2016-2017 Yemen isolates (*n* = 42) belong to sister clusters *Vc*H.9.e and *Vc*H.9.f.

**Figure 2:**
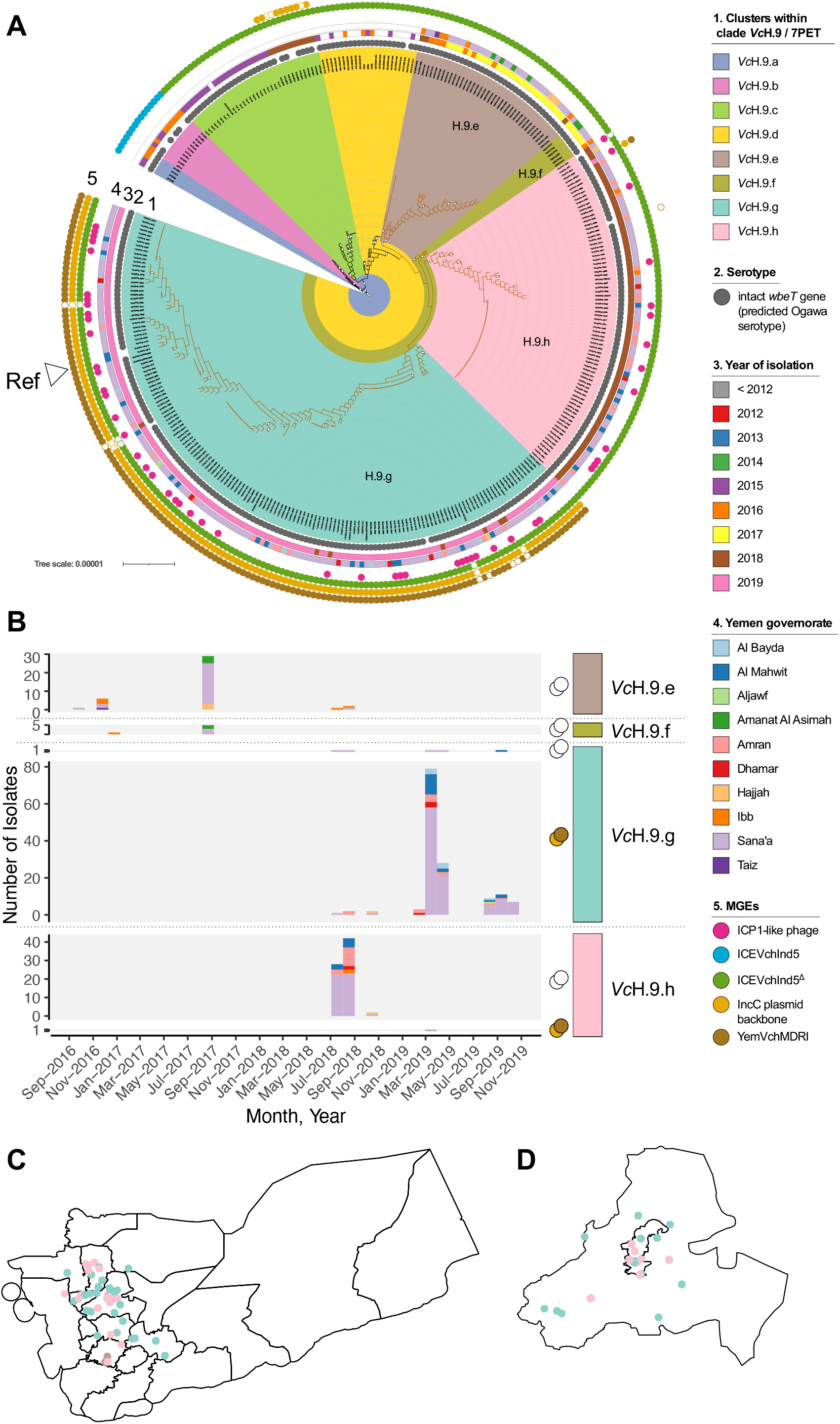
Phylogenetic diversity and spatiotemporal distribution of *Vibrio cholerae* 7PET-T13 isolates (*Vc*H.9) from Yemen. **A**. Subtree of the maximum-likelihood phylogeny of 456 7PET genomes mapped to reference *Vc*H.9 strain CNRVC190243 genome, including 335/456 genomes covering *Vc*H.9 (as defined in Figure S5), which corresponds to the 7PET-T13 sublineage and close South Asian relatives. The full tree containing the 456 genomes is available as supplementary material on Figshare (https://figshare.com/s/4d83a32cce78a52b413e; doi: 10.6084/m9.figshare.16595999) and was obtained based on 2,092 SNP sites from concatenated whole-chromosome alignments. Brown branches indicate the clade grouping all Yemeni 7PET-T13 isolates. Bootstrap support over 70% is indicated by white circles. Phylogenetic clusters within *Vc*H.9 are highlighted with background colours (legend key 1). Coded tracks outside the tree depict the serotype of isolates (ring 2) as predicted from genomic data, year of isolation when isolated in 2012 or later (ring 3), the governorate of isolation if in Yemen (ring 4). The presence of mobile genetic elements (MGEs) is indicated by coloured circles in the outermost track (ring 5): ICP1-like phage (pink), SXT/ICE ICE*Vch*Ind5 (blue), ICE*Vch*Ind5^Δ^ i.e. featuring the characteristic 10-kb deletion in the variable region III (green), IncC plasmid backbone (light brown) and the MDR pseudo-compound transposon Yem*Vch*MDRI (dark brown); filled and unfilled circles indicate different level of coverage in assemblies (see Figure 1 legend). The position of the reference sequence to which all other genomes were mapped to generate the alignment is labelled. The scale bar represents the number of nucleotide substitutions per site. **B**. Frequency of each phylogenetic subcluster among Yemen isolates per month since the onset of the Yemen outbreak. Where relevant, the cluster group is subdivided by the presence or absence of the IncC plasmid as indicated by the filled brown (present) or open (absence) circle on the right of the chart. The contribution of each governorate of isolation is indicated by the coloured portion of each bar. **C** and **D**. A map of Yemen governorates (C) and a focus on the Sana’a and Amanat Al Asimah governorates (inner and outer capital city; D), with dots corresponding to isolates, coloured by phylogenetic subcluster.

### Spatiotemporal distribution of *V. cholerae* isolates

To delineate the evolutionary dynamics of the cholera outbreak in Yemen, we plotted *Vc*H.9 isolates by phylogenetic cluster over time (based on the date of sample collection) and between administrative divisions (linked to reporting hospital). From Figure 2B it is clear that each annual wave was dominated by a single cluster: 2016 and 2017 by *Vc*H.9.e; 2018 by *Vc*H.9.h; 2019 by *Vc*H.9.g. There was no evidence of geographic restriction for any of these clusters, even when accounting for dispersal over time (Fig 2C, 2D; Table S5; Supplemental data online, doi: 10.6084/m9.figshare.19097111). Next, we analysed the relationship between temporal and spatial distances, based on the date and GPS coordinates of sample collection, as well as with the pairwise phylogenetic distances between genomes. We found no significant correlation between the spatial and temporal distances, nor between the spatial and phylogenetic distances (Table S8). These data did show a positive correlation between the temporal and phylogenetic distances (*R*^2^ = 0.181; Mantel test *p*-value < 10^−6^) (Table S8), with root-to-tip distances significantly correlated with sampling date (Pearson’s *R*^2^ = 0.437; *p* < 10^−15^).

We inferred a timed phylogeny for *Vc*H.9 (Figure S6), which revealed that the most recent common ancestor (MRCA) of all Yemeni *V. cholerae* 7PET-T13 genomes was estimated to have existed in February 2015 (95% confidence interval [95%CI], April 2014 and July 2015). Moreover, the MRCAs for clusters *Vc*H.9.e and *Vc*H.9.f (mostly sampled in 2016 and 2017) were dated May and June 2015, respectively, and the MRCAs for clusters *Vc*H.9.g and *Vc*H.9.h (sampled in 2018 and 2019) were dated February and March 2017, respectively. In addition, we dated the MRCA of the clade grouping clusters *Vc*H.9.g and *Vc*H.9.h, which represent the majority of 2018-2019 Yemen isolates, to September 2016 (Figure S6).

The distribution of non-7PET isolates across Yemen was mostly sporadic (Table S4; Suplementary Text; Supplementary data online https://figshare.com/s/73fcd5e1b4958c97ef78, doi: 10.6084/m9.figshare.19097111). However, we characterised a cluster of eighteen closely related *Vc*D isolates belonging to ST555 (Table S6), which we found to differ from each other by 0 to 10 SNPs (average 99.98% ANI similarity). Of these 18 isolates, 13 were isolated over a period of 11 days in late July/early August 2018, two at the end of August, two in October 2018, and one in March 2019 (Table S6). They were obtained from patients in the neighbouring governorates of Sana’a (*n* = 7), Al Mahwit (*n* = 4) and Amran (*n* = 1), which surround the capital city. Genomes from other ST555 isolates, including strains reported as linked to travelers returning to the UK from India in September 2015 and July 2016 (strains 229152 and 338360) ^17^, as well as closest relatives from our core-genome tree, were gathered to build a mapped genome tree of *Vc*D genomes using the complete genome of 2018 Yemen strain CNRVC190247 as a reference (Figure S7). The closest relative to Yemeni ST555 isolates, strain 338360, differs from the *Vc*D ST555 genomes sequenced here by between 763-800 SNPs, ruling out direct clonal relationships.

### Predicted phenotypic properties of *V. cholerae* isolates

Consistent with our previous report^7^, Yemeni *Vc*H.9 isolates – which all belong to 7PET-T13 sublineage – all carried genes or mutations known to confer resistance to trimethoprim (*dfrA1*) and to nalidixic acid (*gyrA*_S83I and *parC*_S85L). They also carried the *Vibrio* pathogenicity island 1 (VPI-1, encoding the toxin co-regulated pilus TCP), VPI-2, the *Vibrio* seventh pandemic islands I and II (VSP-I and VSP-II), and the CTX prophage, which all featured the cholera toxin genes, *ctxAB*, of the allelic type *ctxB7*. None of the non-7PET genomes from Yemen possessed a CTX prophage or the *ctxAB* genes. However, Yemni isolates belonging to *Vc*K (ST170, related to previously described lineage MX-2), which were derived from the stool of patients presenting cholera-like disease, carried all the genes coding for the TCP.

These *ctxAB*^-^, *tcpA*^+^ *Vc*K genomes also carried the O1 LPS O-antigen biosynthetic gene cluster, consistent with what has been seen previously in related non-7PET isolates^18^. The genomes of the *Vc*D isolates belonging to ST555, ST1020, ST1498 and the *V. paracholerae* isolate (ST1499), carried LPS O-antigen biosynthetic gene clusters encoding unknown serogroups; these were conserved within and specific to each ST (Table S4; Figure S7A). Of the 216 Yemeni 2018-2019 *Vc*H isolates, 213 were predicted to produce a serogroup O1 LPS O-antigen based on presence of a full biosynthetic gene cluster; in the three remaining assemblies this genomic region was interrupted (YE-NCPHL-19012) or completely missing (YE-NCPHL-18033 and YE-NCPHL-19140), likely due to limited genome sequence coverage (Table S3). All predicted O1 serogroup isolates were predicted to be Ogawa serotype except two that showed a disruption in *wbeT*, indicative of an Inaba phenotype (YE-NCPHL-18053 and YE-NCPHL-19014, with gene truncation and point mutation respectively; Table S5). These predictions were imperfectly reflected by the results of serological assays conducted at NCPHL (Table S2; Figure S8), suggesting issues in initial laboratory testing (see Supplementary Text).

### Genome variation of *Vc*H.9 (7PET-T13) isolates circulating in Yemen

Given the change in antimicrobial suceptibility seen in the 2018-2019 Yemen isolates, we compared in detail all of the *Vc*H.9 isolate genomes from Yemen to each other and related isolates taken elsewhere, focusing on genotypic traits that were conserved in pandemic sublineages occurring in Yemen. We identified three, four, and 21 fixed SNPs in the the crown clade containing *Vc*H.9.e,f,g,h, the clade containing *Vc*H.9.g,h, and *Vc*H.9.h, respectively (including 2, 2 and 11 non-synonymous SNPs, respectively; Table S9). Changes fell largely within genes predicted to be involved in carbohydrate metabolism, signal transduction and chemotaxis, none of which could be directly linked to change in virulence (Table S9).

Previously, the 2016-2017 Yemeni isolates carried an SXT ICE differing by only three or four SNPs from the ICE*Vch*Ind5/ICE*Vch*Ban5 reference sequence (Genbank accession GQ463142.1)^19^, but which possessed a 10-kb deletion in variable region III, which explained the phenotypic loss of resistance to streptomycin, chloramphenicol and sulphonamides (only retaining resistance to trimethoprim via the *dfrA1* gene)^7^. All 2018-2019 *Vc*H.9 genomes carried the same SXT ICE variant, with a maximum of 2 SNP differences and displaying the same deletion. Hence, the change in antimicrobial resistance profile was not linked to variation in SXT ICE.

Looking across all genes within the pangenome, the only variation directly associated with the Yemen 2018-2019 genomes, compared to those sequenced from 2016-2017, was the presence of a novel 139-kb plasmid, which we named pCNRVC190243 (Table S10). The backbone of this new plasmid includes a replicon of the IncC type, as well as genes encoding a complete type F conjugative apparatus and a MOBH-type relaxase, suggesting it is self-transmissible. Plasmid pCNRVC190243 also carries a 20-kb genomic region (which we denoted Yem*Vch*MDRI) predicted to encode a quaternary ammonium compound efflux pump (*qac*), an extended-spectrum beta-lactamase (ESBL; *bla*PER-7), sulphonamide resistance (*sul1*), aminoglycoside resistance (*aadA2*), and macrolide resistance (*mph*(A), *mph*(E) and *msr*(E)) (Figure 3; Table S4). Yem*Vch*MDRI is a pseudo-compound transposon (PCT) – a structure bounded by IS*26* elements^20^ – and includes a class 1 integron with *aadA2* encoding resistance to streptomycin and spectinomycin as a gene cassette, associated with an IS*CR*1 element carrying the ESBL *bla*PER-7 gene, a structure similar to one previously seen in *Acinetobacter baumanii*^21,22^. We found that pCNRVC190243 was present in 6/89 (6.7%) Yemeni *Vc*H.9 isolates from 2018, but this rose to 100% (151/151) in 2019 (Figure 2B). This was linked to phylogenetic cluster, with only 1/79 (1.3%) *Vc*H.9.h isolates harbouring the plasmid, compared to all (156/156) *Vc*H.9.g isolates (Figure 2A).

**Figure 3:**
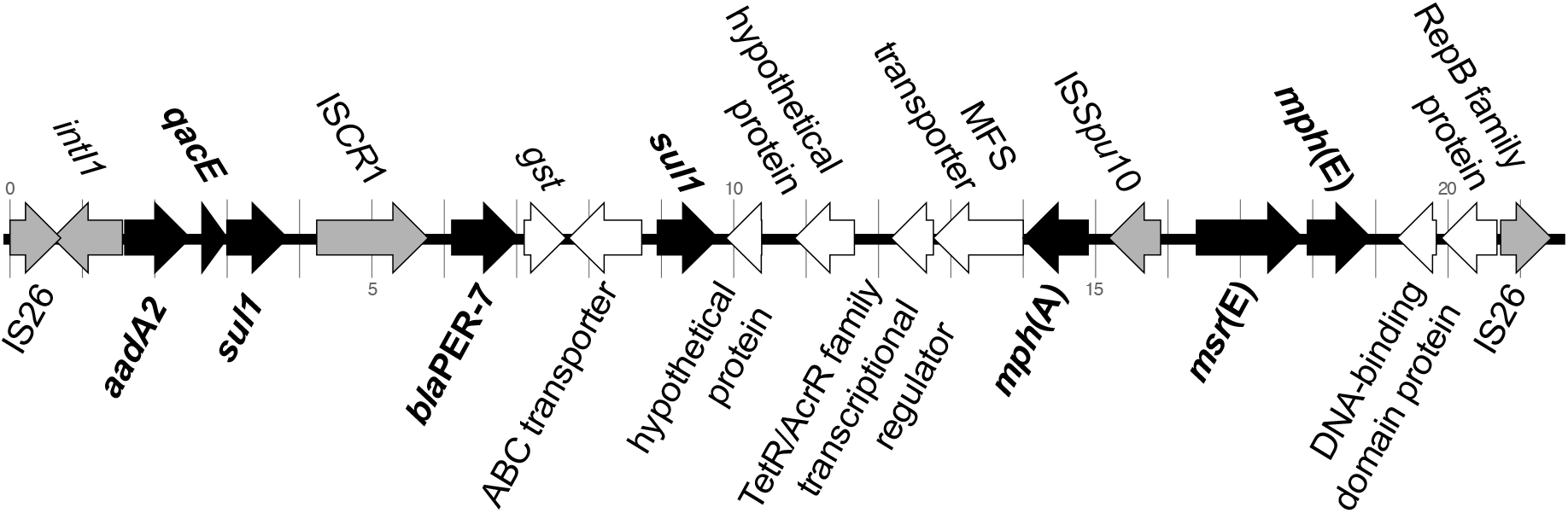
Genetic organisation of the MDR pseudo-compound transposon Yem*Vch*MDRI. Antimicrobial resistance (AMR) genes are filled in black and labelled in boldface; genes encoding endonucleases transposases and other genes involved in genetic mobility are filled in grey. Genomic position is indicated by tickmarks every kilobase, in reference to the pCNRVC190243 plasmid coordinates.

### Distribution and relatedness of MDR mobile genetic elements

Analysis of the broader phylogenetic context of pCNRVC190243 suggested at least three independent acquisitions of this plasmid (and associated Yem*Vch*MDRI), since it was also present in three *Vc*D (ST1499 and ST1020) and two *Vc*K (ST170) isolates collected in 2019 in Yemen. Comparing the full-length sequence of all pCNRVC190243 plasmids from *Vc*H.9 and *Vc*K and *Vc*D isolates showed all sequences were identical except for two isolates: one varied by a single SNP resulting in an amino-acid change S71F in the sulphonamide resistance protein Sul1 (YE-NCPHL-19105; G26720A SNP); the other by a single intergenic SNP. We also found the Yem*Vch*MDRI element integrated into chromosome 2, without the pCNRVC190243 backbone (Figure 1; Supplementary Text) in all eighteen of the ST555 isolates.

Searching a broader prokaryotic genome database, closely related but non-identical elements were found in different combinations in other *V. cholerae* and diverse bacterial taxa: an IncC plasmid, named pYA00120881 (GenBank accession MT151380), was identified in 13 closely related *Vc*H.9.a and *Vc*H.9.c isolates (Figure 2A) that were collected in 2015 and 2018 in Zimbabwe^15^. The backbones of these IncC plasmids share 99.98% nucleotide sequence identity, but pYA00120881 carries a different MDR genomic region – featuring a *bla* gene encoding a CTX-M-15 ESBL – inserted at the same locus (Figure S9). Furthermore, 59 *V. cholerae* O139 (ST69) isolates collected in China from 1998 to 2009^23,24^ (unpublished genomic data released in BioProject PRJNA303115; Table S11) carry IncC-type plasmids that show similarity to pCNRVC190243 and also include Yem*Vch*MDRI-like PCT elements, albeit lacking IS*CR*1 and its associated *bla*_PER-7_ gene.

Importantly, when using the Yem*Vch*MDRI sequence alone for database searches, we found the genome of *V. cholerae* ST555 strain 338360 (Table S6) shared 100% nucleotide identity with the complete Yemeni ST555 Yem*Vch*MDRI sequence, including the *bla*_PER-7_-carrying IS*CR*1 (IS*CR*1*bla*_PER-7_; Table S12). Likewise, IS*CR*1*bla*_PER-7_ has also been previously observed in the genomes of *A. baumanii* strains^21,25^ from France and the United Arab Emirates (UAE). Those from UAE were located on the plasmid pAB154, where the sequence homology with IS*CR*1*bla*_PER-7_ extended beyond the canonical element and included Yem*Vch*MDRI flanking regions, suggesting that the IS*CR*1*bla*_PER-7_ carried by pAB154 is derived from Yem*Vch*MDRI, or a closely related element (Figure S10). Moreover, outside of *V. cholerae*, pCNRVC190243- and/or Yem*Vch*MDRI-like elements are widely distributed with *Escherichia coli, Salmonella enterica* and *Klebsiella pneumoniae* genomes presenting >95% shared nucleotide *k*-mers (Table S11, S12; Figure S11), with the closest matches outside of *V. cholerae* being seen in *K. pneumoniae*. This indicates that similar regions may be widely distributed in MGEs across bacterial taxa.

A recent study reported that two anti-plasmid defence systems, DdmABC and DdmDE, cause the instability of plasmids in *V. cholerae* cells, including those with IncC-type replicons^26^. These proteins are encoded by all 7PET genomes, which according to this study would explain the incapacity of plasmids to be maintained in a 7PET background. We here show that a 139-kb IncC-type plasmid has been stably propagated in a clone of the 7PET lineage. We verified the presence and integrity of the DdmABC and DdmDE systems and found they were present and intact in all 7PET genomes in our 882 assembled genome dataset – including those harbouring pCNRVC190243 (Table S5). This shows that these defence systems are not sufficient to destabilise pCNRVC190243 to the point of it being lost from the population within the 15-month period covered by our study, or even for the following two years, as suggested by 2021 antibiotic susceptibility profiles.

### Discussion

Characterising the genomic nature of the pathogens causing an outbreak can reveal changing epidemiological dynamics through adaptative evolution of the pathogen or the introduction of distinct pathogen lineages. Such events may lead to the emergence of a more virulent or drug resistant genotype of the pathogen, and impact disease control efforts. Our genomic epidemiology analysis shows that despite seasonal fluctuation, the vast majority of cholera in Yemen is caused by the 7PET-T13 lineage (*Vc*H.9), and is derived from a single introduction into Yemen. Using a larger sample set, we refined our previous date estimate^7^ and show that the progenitor of the Yemeni outbreak emerged between April 2014 and September 2015, contemporarily to the onset of the civil war in Yemen, and existed one to two years prior to the declaration of a cholera outbreak^3,7^.

Using our high-resolution phylogenomic tree, we were able to subtype the majority of Yemeni genomes into four different phylogenetic clusters that dominated the outbreak at different points in time. We observed two large clonal expansions for the sister clades that dominated 2018 and 2019 (clusters *Vc*H.9.h and *Vc*H.9.g, respectively), which both emerged in early 2017. Founding effects at the onset of each cholera season, associated with rapid expansions, may explain the dominance of each cluster in these respective epidemic waves. However, it is also possible that an adaptive advantage participated in driving the replacement of *Vc*H.9.h by its sister clone, *Vc*H.9.g. In the absence of samples from subsequent years (with surveillance efforts hampered by the Covid-19 pandemic), it was not possible to establish whether these or another MDR lineage persisted past 2019.

In Yemen, pregnant women and children (one third of cholera patients were aged 15 or under; Table S7; ref. 11) were treated with erythromycin and azithromycin between 2016 and late 2018. The 2019 wave of the Yemen cholera outbreak was associated with a sudden change in antibiotic resistance profile, from being largely sensitive to antimicrobials between 2016-2018, to being resistant to multiple therapeutically relevant drugs in 2019. Our data showed that this phenomenon coincided with the appearance in late 2018 of plasmid pCNRVC190243 in isolates belonging to *Vc*H.9.g, the phylogenetic cluster which dominated our 2019 samples. Plasmid pCNRVC190243 carries the pseudo-compound transposon Yem*Vch*MDRI, which in turn includes a type 1 integron and the IS*CR*1*bla*_PER-7_ element. These elements confer resistance to third-generation cephalosporins, aminoglycosides, macrolides and sulphonamides (and, combined with the *dfrA1* gene present on the SXT ICE, to co-trimoxazole), plus disinfectant tolerance provided by the *qac* gene^27^. The acquisition of Yem*Vch*MDRI element by an ancestor of *Vc*H.9.g was followed by its dramatic spread – a clonal expansion which we show to occur in 2018 (Figure S6), a time when there would have been a selective pressure towards macrolide resistance in symptomatic cases due to the large-scale administration of these drugs.

We also identified a small number of non-7PET *V. cholerae* amongst 2018-2019 Yemen isolates: 8% of unique clinical isolates (21/254) and 30% of environmental isolates (3/10) belonged to three diverse lineages. The location and times of isolation of these non-7PET *V. cholerae* isolates suggest they largely represented sporadic infection events linked to endemic strains. The only sizable cluster of non-7PET isolates were the 18 *Vc*D/ST555 isolates which had near-identical genomes and isolated in 2018 and 2019 in several districts near the capital city of Sana’a (Figure 1; Figure S7; Table S4). The short time range in which 15 of these strains were isolated (31 days in July-August 2018) could be explained by repeated acquisitions from a point source, although we cannot rule out that they stem from small-scale outbreaks, as has been reported previously for non-O1/non-O139 strains^28,29^. However, we found no evidence of long range spread of these non-pandemic clones across Yemen, characteristic of 7PET *V. cholerae* isolates linked to epidemic disease. Importantly, the reappearance of this *Vc*D/ST555 genotype later in October 2018 and March 2019, with as little as 2 SNPs difference from summer 2018 isolates, could suggest this genotype is able to persist in the environment, possibly through similar ecological mechanisms as those that lead to the seasonal dynamics of epidemic cholera following its initial introduction^11^. These ST555 strains might in fact represent an endemic population of *V. cholerae* that can be carried without causing any disease. These ST555 strains could have been isolated from a gut co-colonised by a cholera-causing 7PET strain, in an epidemic context where the pathogen is routinely isolated from cholera patients using culture and enrichment techniques that are selective of the whole *V. cholerae* species. This hypothesis of incidental isolation of ST555 strains in samples also containing a toxigenic 7PET strain is supported by the original serotyping of all samples as O1 (Table S1; Supplementary Text), and the positive detection of the *rfbO1* marker by PCR in samples from which the four ST555 strains were isolated at the Institut Pasteur (IP) – samples which, when sequenced at the Wellcome Sanger Institute (WSI), yielded 7PET genomes (Figure S3; Table S13; Supplementary Text).

Sequences completely or near identical to plasmid pCNRVC190243, carrying the PCT Yem*Vch*MDRI, were present in i) 7PET-T13 (*Vc*H.9) isolates, ii) all isolates from two of the three different STs of *Vc*D (ST1499 and ST1020), and iii) the two *Vc*K/ST170 isolates; all of which were collected in 2019 in Yemen. The only *Vc*H.9.h isolate from 2019 also carried this plasmid. It is possible to explain these observations by multiple acquisitions of the plasmid from independent sources or, more parsimoniously, as direct horizontal gene transfer events between the *V. cholerae* lineages we report here. The large population sizes attained by the epidemic lineages in Yemen make the latter hypothesis more likely, in a scenario of spill over from the dominant cluster at the time, *Vc*H.9.g. However, we could not infer directionality due to the limited available sampling from the diverse lineages in this study.

The Yem*Vch*MDRI PCT was also carried chromosomally by Yemeni ST555 strains. This PCT is itself a composite element, including IS*CR1bla*_PER-7_, a rare element that has only been observed once in another *V. cholerae* background – the relatively closely-related ST555 strain 338360 (436 SNPs vs. reference strain CNRVC019247), isolated from a traveler returning from India – and in two plasmids associated with *A. baumanii* strains isolated in Gulf countries. Comparison of these homologous IS*CR1* elements suggests they all are derived from the same ancestral element (Figure S10). Presence of the full, identical PCT Yem*Vch*MDRI in two closely related, but distinct ST555 strains isolated from completely different geographical origins, suggests this element is stably associated with this genotype. From this, we can speculate that Yem*Vch*MDRI was originally present in Yemen in a ST555 genomic background, and later combined with an IncC plasmid backbone to produce pCNRVC190243. Again, directionality cannot be confidently inferred because of uneven sampling of host lineages, and the potentially large number of unobserved donor bacteria.

Whilst pCNRVC190243 is a novel element, plasmids such as pYAM00120881 identified in *Vc*H.9 *V. cholerae* from Zimbabwe in 2015 and 2018^15^ shared almost identical plasmid backbones. In addition, similar plasmids, some of which also carry Yem*Vch*MDRI-related elements, have been observed in *V. cholerae* O139 isolates from China as well as detected in a range of other bacterial genera, illustrating how widely distributed these IncC plasmids are. Similarly, Yem*Vch*MDRI may occur in more diverse and more widely spread genomic backgrounds that haven’t been sampled yet. It is therefore possible that parts of this plasmid have combined outside of Yemen from identical or similar genomic sources, independently from the Yem*Vch*MDRI-carrying Yemeni ST555 strain.

While other MDR IncC plasmids were previously observed in *V. cholerae* in DRC, Kenya (in a T10 sublineage genetic background) and Zimbabwe (T11 background in 2015), these were only linked to sporadic cases or small-scale cholera outbreaks^30^, despite selective conditions linked to the widespread and uncontrolled use of antibiotics. Recently, it has been shown that two defence systems, called DdmABC and DdmDE had the capacity to destabilise plasmids, including large IncC-type plasmids^26^. These defence systems, encoded in all 7PET *V. cholerae* genomes, were proposed to be responsible for the lack of maintenance of MDR plasmids in populations of this pandemic lineage when not under stringent selective pressure for antibiotic resistance. A first exception to the pattern of plasmid instability in 7PET *V. cholerae* was the Zimbabwean cholera outbreak of 2018, which lasted six months and produced over 10,000 suspected cases, and was associated with a strain of the T13 background carrying the MDR IncC plasmid pYA00120881^15^. The Yemeni cholera outbreak provides a further example, with the T13 strain of the *Vc*H.9.g clone, carrying pCNRVC190243, being presumably associated with more than a million suspected cases recorded since its emergence in late 2018. However, the intact presence of the genes encoding the Ddm proteins in Yemeni and Zimbabwean *Vc*H.9/7PET-T13 genomes presenting MDR IncC plasmids indicates there may be other mechanisms that impact plasmid stability in 7PET genomes.

One possibility would be that an unknown environmental factor has applied a consistently strong selective pressure for a trait carried by these plasmids. Even though the treatment of cholera patients with macrolides was stopped in Yemen in early 2019, antibiotic pressure remains a potential selective factor, as antibiotics and particularly azithromycin have been reported to be overused by the general population in Yemen during the Covid-19 crisis^31^. Another possible factor would be the interaction with other mobile elements, including ICP1 phages, which we detected in a significant fraction of the samples (see Supplementary Text). It has also been proposed previously that the presence of an SXT ICE in these genomes could prevent the stable replication of an IncC-type plasmid, through an unknown functional interference mechanism^7,15^. The unique occurrence of a 10-kb deletion in the SXT ICE (ICE*Vch*Ind5/ICE*Vch*Ban5) in T13 isolates may provide these genomes with the novel capacity to stably host an IncC plasmid; molecular genetic investigation of this locus should be conducted to test whether it encodes another plasmid destabilisation factor. Whatever the mechanism, it appears that both MDR elements SXT ICE and IncC plasmid are stably propagated together in the Yemeni T13 strain, which population in Yemen has reached a unprecedented size. This emerging MDR strain has therefore a high potential to spread and seed further adapted lineages, as well as to disseminate its MDR plasmid and PCT to other organisms.

## Conclusion

The emergence of this multi-drug resistant pathogen demonstrates the necessity of continued genomic surveillance of the microbial population associated with the ongoing Yemen cholera outbreak, and for new outbreaks that may take place in regionally connected areas. Such surveillance will enable Yemeni public health authorities to rapidly adapt clinical practices to minimize AMR selective pressures. This also warrants increased efforts in research on the molecular mechanisms and evolution of interactions between mobile genetic elements, to learn about the constraints ruling their colonization of bacterial genomes. Such knowledge is essential for us to be able to disentangle the role of MGEs from that of their bacterial hosts in driving epidemics, so to propose practical definitions of pathogens that focus on the relevant genes, mobile elements or prokaryotic organisms, and to implement appropriate molecular epidemiology surveillance schemes.

## Materials and Methods

### Definitions and surveillance data

Cholera cases were notified to the the Ministry of Public Health and Population of Yemen (MPHP) and recoded through the Electronic Disease Early Warning System (eDEWS)^2^. Suspected and confirmed cholera cases were defined according to the WHO in a declared outbreak setting. Briefly, a suspected case is any person presenting with or dying from acute watery diarrhoea (AWD) and a confirmed case is a suspected case with *Vibrio cholerae* O1 or O139 infection confirmed by culture.

### Sample collection, microbiological testing and clinical metadata

Clinical samples, i.e. stool and rectal swabs, were collected in Yemen by epidemiological surveillance teams from suspected cholera cases during 2018 and 2019^11^ and were transported to the National Centre of Public Health Laboratories (NCPHL) in the capital city Sana’a in Cary-Blair transport medium (Oxoid, USA). To probe the diversity of vibrios shed by unreported cholera cases, as well as *V. cholerae* that may naturally occur in effluent waters, environmental samples were collected during the day time in October 2019 from the sewage system around Sana’a city and the vicinity and then transported to NCPHL for testing; each sample was collected in sterile bottles containing enrichment media comprised of 250 mL of sewage and alkaline peptone broth (APB, Difco Laboratories, Detroit, Michigan) at a 1:1 ratio and incubated for 20 h at room temperature including the transportation time into the NCPHL and processed as described previously^32^. All samples were cultured and identified according to the Centers for Disease Control and Prevention (CDC) guidelines^33^. Resistance to antibiotics was tested by the disk diffusion method according to the CLSI guidelines^34^ for a range of antibiotics as described in Table S1.

Live clinical isolates (n=120) were sent to the Institut Pasteur (IP; Paris, France), where only 21 samples were culture positive, due to poor sample preservation during shipment (Table S2; Figure S3), leading to the final isolation of 22 *V. cholerae* strains (including two from mixed culture YE-NCPHL-18020). Strains re-isolated at IP were characterized by biochemical and serotyping methods according to standard practice of the French National Reference Centre for Vibrios and Cholera (CNRVC)^35^. Separate antibiotic susceptibility testing (Table S2) was performed by the disk diffusion method according to EUCAST guidelines (EUCAST 2020^36^) and MIC determination using the Sensititre™ (Thermo Scientific) and the Etest ® (bioMérieux, Marcy-l’Étoile, France) systems. Interpretation into S (Susceptible), I (Intermediate), and R (Resistant) categories was performed according to the 2020 edition of EUCAST recommendation on interpretation of the diameter of the zones of inhibition of *Enterobacteriaceae*^37^, and to the 2013 CA-SFM (Comité de l’Antibiogramme de la Société Française de Microbiologie) standards for *Enterobacteriaceae*^38^ for antibiotics for which critical diameters are no longer reported in the latest published guidelines. *E. coli* CIP 76.24 (= ATCC 25922) was used as a reference strain.

### DNA extraction and sequencing

Genomic DNA was extracted at the NCPHL from subcultures inoculated with single bacterial colonies and grown in nutrient agar (Oxoid, USA) at 37°C overnight according to the manufacturer instructions (Wizard® Genomic DNA Purification kit, Promega, UK). Genomic DNA samples (derived from 10 environmental and 250 clinical samples, which includes the 120 samples sent to IP) were sent to the Wellcome Sanger Institute (WSI; Hinxton, UK) and sequenced on the WSI sequencing pipeline (Figure S3) using the Illumina HiSeq platform X10 as previously described^28^.

Two MDR *V. cholerae* strains were selected among the 22 held at the IP for long-read sequencing. The first strain, CNRVC190243 (= YE-NCPHL-19014-PI), a 7PET *V. cholerae* O1 strain was sequenced by Single-Molecule Real-Time (SMRT) sequencing (Pacific Bioscience). The genomic DNA was prepared at the IP as follows: strain CNRVC190243 was cultured in Brain-Heart-Infusion (BHI) broth (Difco) overnight at 37 °C with shaking (200 rpm—Thermo Scientific MaxQ 6800). Then, 100 µL of the overnight culture was inoculated into a 10 ml BHI broth and cultured 2 hours at 37°C with shaking. After centrifugation, the bacterial cells were processed as described previously^39^, except that MaXtract High Density columns (Qiagen) were used (instead of phase lock tubes) and DNA was resuspended in molecular biology grade water (instead of 10 mM Tris pH 8.0). Library preparation and the sequencing were performed by the GATC platform (Eurofins Genomics Europe Sequencing GmbH; Konstanz, Germany) using their standard genomic library protocol and PacBio RS sequencer. The second strain, CNRVC190247 (= YE-NCPHL-18035-PI), a non-O1/non-O139 *V. cholerae* strain further characterized as ST 555, was sequenced using the MinION nanopore sequencer (Oxford Nanopore Technologies). Genomic DNA was prepared at the IP as follows: strain CNRVC19247 was cultured in alkaline nutrient agar (casein meat peptone E2 from Organotechnie, 20 g; sodium chloride from Sigma, 5 g; Bacto agar from Difco, 15g; distilled water to 1 L; adjusted to pH 8.4; autoclavated at 121°C for 15 min) overnight at 37 °C. A few isolated colonies of the overnight culture were inoculated into a 20 ml of Brain-Heart-Infusion (BHI) broth and cultured until a final OD_600_=0.8 at 37°C with shaking (200 rpm). After centrifugation, the bacterial cells were processed as described above. The library was prepared according to the instructions of the “Native barcoding genomic DNA (with EXP-NBD104, EXP-NBD114, and SQK-LSK109)” procedure provided by Oxford Nanopore Technology. The sequencing was then performed using a R9.4.1 flow cell on the MinION Mk1C apparatus (Oxford Nanopore Technologies). The genomes of 21/22 strains cultivated at the IP (all but CNRVC190251, which was isolated later; Table S2) were also sequenced using Illumina short-read technology at the IP using the equipment and services of the iGenSeq platform at the Institut du cerveau et de la moëlle épinière (Paris, France) from genomic DNA extracted with the Maxwell 16-cell DNA purification kit (Promega) in accordance with the manufacturer’s recommendations.

### Genome assembly and annotation

The 260 sequencing read sets produced at the WSI (Figure S3) were processed with the WSI Pathogen Informatics pipeline^40^: quality of sequencing runs was assessed based on quality of mapping of 10% reads to the genome of reference strain N16961 (GenBank Assembly accession GCA_900205735.1) using the Burrows-Wheeler Aligner (BWA)^41^; read sets passed the check if at least 80% bases were mapped after clipping, the base and indel error rate were smaller than 0.02, and less than 80% of the insert sizes fell within 25% of the most frequent size. Contamination was assessed manually based on Kraken classification of reads using the standard WSI Pathogen reference database, which contains all viral, archaeal and bacterial genomes and the mouse and human reference published in the RefSeq database as of the 21^st^ May 2015 (Table S3). Sequences were assembled *de novo* into contigs as described previously^42^, using SPAdes v3.10.0 as the core assembler^43^. Poor assemblies were filtered out if differing of more than 20% from the expected genome size of 4.2 Mb, or when more than 10% of reads were assigned by Kraken to another organism than *V. cholerae* (notably including the *Vibrio* phage ICP1) or to synthetic constructs, or were unclassified. This led to the omission of 28 genome assemblies, resulting in 232 high-quality assembled genomes. The genome of strains CNRVC190243 and CNRVC190247 were assembled based on long and short reads using a hybrid approach with UniCycler^44^ v0.4.7 and v0.4.8, respectively, using pilon^45^ v1.23 for the polishing step, to produce high-quality reference sequences comprised of both chromosomes and, for strain CNRVC190243, of an additional plasmid, pCNRVC190243. New genomes were annotated with Prokka version v1.5.0^46^.

### Contextual genomic data (882 “assembled *V. cholerae* genomes” dataset)

To provide phylogenetic context, we also included in this analysis previously published genome sequences from a globally representative set of isolates. We first gathered genome assemblies generated at the WSI using the pipeline described above based on previously published short reads sets from *V. cholerae* isolates belonging to sublineage T13 of 7PET Wave 3 (7PET-T13) and from strains isolated in close spatio-temporal context i.e. within a decade in Africa and South Asia (where the ancestor of T13 is thought to originate^7^). These include all 42 Yemen 2016-2017 isolates^7^, 103 recent isolates from East Africa including from Kenya^7^, Tanzania^47^, Uganda^48^ and Zimbabwe^15^ and 74 isolates from South Asia ^49^. In addition, we included genomes spanning the wider diversity of *V. cholerae*, including all 119 genomes from China^18^, as well as 312 genomes from the collections of contextual genomes used in previous studies^7,28^. Together with the 232 Yemen 2018-2019 isolate genome assemblies (see above), our final dataset consisted of 882 assembled *V. cholerae* genomes (Table S4; Figure S3).

### Identification and typing of mobile genetic elements, virulence factors, AMR genes and anti-phage defense systems

The presence of AMR genes, plasmid replicon regions or virulence factors were predicted using Abricate^50^, searching the reference databases NCBI AMR+^51^, Plasmidfinder^52^ or VFDB^53^, respectively. BLASTN^54^ (v2.7.1+, with default parameters) was used to identify known mobile genetic elements (MGEs): the SXT/ICE ICE*Vch*Ind5 (GenBank accession GQ463142.1); ICP1-like vibriophages ICP1_VMJ710 and ICP1_2012_A (GenBank accessions MN402506.2 and MH310936.1, respectively)^55^ and the ICP1-like vibriophage YE-NCPHL-19021, which genome was the only assembled contig from the reads obtained from sample YE-NCPHL-19021 (this study; Genbank accession MW911613.1); the IncC-type plasmid pCNRVC190243, obtained from the hybrid assembly of strain CNRVC190243 described above (this study; ENA sequence accession OW443149.1); the MDR pseudo-compound transposon (PCT) Yem*Vch*MDRI, extracted from this plasmid (positions 16,442 to 36,862); PICI-like elements (PLE) 1, 2 and 3 (GenBank accessions KC152960.1, KC152961.1, MF176135.1)^56,57^. Absence of elements was verified at the read level as described below. Sequences similar to the reference sequences of the plasmid pCNRVC190243, the MDR PCT Yem*Vch*MDRI and the ICP1-like phage genome YE-NCPHL-19021 were also searched in a database of 661,405 genome assemblies ^58^ using a *k*-mer-based COBS index ^59^; alignment of best matches were further characterized using BLASTN. We typed the conjugation apparatus of pCNRVC190243 with CONJScan ^60^ on the Pasteur Institute Galaxy server (Galaxy Version 1.0.5+galaxy0). We searched for presence of CRISPR-Cas arrays using MacSyFinder^61^ v1.0.5 with default parameters and the built-in Cas system reference database; genomes positive for Cas systems were further analysed with CRISPRCasFinder ^62^ on the Pasteur Institute Galaxy server to retrieve CRISPR arrays.

### Prediction of serotype, serogroup and multi-locus sequene type

To predict the antigenic serogroup, we screened the assemblies against a custom reference database using Hamburger^63^. In brief, a database was constructed by selecting flanking and marker genes for the operon encoding the *V. cholerae* O-antigen, with representative genes for both O1 and non-O1 serogroups included (Table S15). Gene sequences were individually aligned using Clustal Omega (version 1.2.4), prior to HMM construction with HMMER (version 3.2.1) and concatenation of the HMM alignments. Assemblies were screened against this database using Hamburger (version 836a77c)^64^ to identify the operon, and genetic structure was compared across the assemblies and references to designate serogroups. The HMMER database is available online at https://figshare.com/s/5dd21a52f0d5a39a670f (doi: 10.6084/m9.figshare.19575148).

For O1 serotype prediction (Inaba or Ogawa), we used a combination of approaches including BLASTN search against the 882 assembled *V. cholerae* genomes (as described above) and ARIBA (v2.14.6+, with default parameters)^65^ to screen the sequencing read sets against the *wbeT* gene sequence from strain NCTC 9420 (positions 311,049-311,909 of GenBank accession CP013319.1) as a reference, as previously described^28^. Multi-locus sequence typing (MLST) of non-7PET isolates was conducted on PubMLST.org^66^ under the non-O1/non-O139 *V. cholerae* seven-gene typing scheme.

### Identification of single nucleotide variants (456 “mapped 7PET genomes” and 33 “mapped *Vc*D genomes” datasets)

For variant calling, Illumina short reads were mapped against the novel reference genomes from strains CNRVC190243 and CNRVC190247, or the in-house MGE database described above. We mapped all 260 short read sets from 2018-2019 Yemeni isolates sequenced at the WSI, including those 28 read sets which assembly showed low coverage or appeared contaminated with phage genomes (Table S3), so to recover variation data evidenced at the read level, provided reads were mapped at a sufficient depth (see below). We also mapped read sets from the 21 strains sequenced at the IP, and from contextual isolates of the 7PET-T13 sublineage and close relatives (see “Contextual genomic data”), for a total of 468 mapped genomes. Reads were trimmed with Trimmomatic, mapped to both CNRVC190243 reference chromosomes with BWA-MEM and the IncC plasmid pCNRVC190243. Mapped genomes with an average read depth below 5x over the two chromosomes were deemed of insufficient read depth and were excluded (12 read sets mapped to CNRVC190243, all from this study and generated at WSI, were excluded for a final set of 456 mapped 7PET genomes [Table S5]; no read set mapped to CNRVC190247 was excluded). We used the software suite samtools/bcftools^67^ v1.9 to call single nucleotide variants with a minimum coverage of 10x read depth; see custom script ‘map_yemen_reads2MGEs.sh’^68^ for a detailed description of the parameters used. Resulting consensus sequences were combined into a whole-genome alignment, which was processed with snp-sites ^69^ to produce a single nucleotide polymorphism (SNP) alignment.

Overall genome similarity was assessed by computing SNP distances based on the above alignments using the function ‘dist.dna’ from the R package ‘ape’^70^, and average nucleotide identity (ANI, accounting for unaligned regions) was computed using fastANI^71^ v1.3 with default parameters.

### Phylogenetic inference

The Pantagruel pipeline^72^ was used to infer a maximum-likelihood (ML) “core-genome tree” using the “-S” option and otherwise default parameters. Briefly, 291 single-copy core-genome genes (with expected high degree of sequence conservation and relatively low prevalence of HGT compared to other core genes) were extracted from the 882 assembled *V. cholerae* genomes, their alignments were concatenated and the resulting supermatrix was reduced to its 37,170 polymorphic positions, from which a ML tree was computed from RAxML v8.2.11^73^ (model ASC_GTRGAMMAX using Stamatakis’ ascertainment bias correction; one starting parsimony tree; 200 rapid bootstraps for estimating branch supports); supporting supplementary data are available on Figshare at https://figshare.com/s/3fe31c131b00a2a08bb9. Phylogenies were also inferred from whole-genome alignments of the concatenated consensus sequences of both chromosomes from the SNP alignment of the 456 mapped 7PET genomes and 33 mapped *Vc*D genomes. These alignments contained 2,092 and 91,312 polymorphic positions, respectively, and were used as input to RAxML-NG v1.0.1^74^ to build the ML “mapped genome trees” using the following options: “all --tree pars{10} --bs-trees 200 --model GTR+G4+ASC_STAM”. Alternative topologies were compared using RAxML-NG option “--sitelh” to generate per-site likelihood values and the ‘SH.test’ function from the ‘phangorn’ R package^75^ to test hypotheses.

The 882 assembled *V. cholerae* core-genome tree was rooted using the clade of sequences identified as *V. paracholerae*^14^ as an outgroup. The remaining part of the tree (*V. cholerae sensu stricto*) was subdivided into clades named *Vc*A to *Vc*K based on visual examination with the aim to coincide with previously described lineages such as 7PET, Gulf Coast, etc. or based on balanced subdivisions of the tree diversity. *Vc*H, corresponding to the 7PET lineage, was further subdivided into clades of even depth, named Subclades H.1 to H.9. The 456 mapped 7PET genomes were similarly classified into clusters based on the tree topology, with genomes assigned to subclades named *Vc*H.5, *Vc*H.6, *Vc*H.8 or *Vc*H.9 (according to their position in the 882 assembled *V. cholerae* core-genome tree). Genomes belonging to *Vc*H.9, which corresponds to the 7PET-T13 sublineage, were further separated into *Vc*H.9.a to *Vc*H.9.h, based on visual examination of the tree structure and aiming to maximise uniformity of the spatio-temporal metadata associated to genomes in each cluster; clusters correspond to clades, either entirely or at the exclusion of another cluster included in the clade i.e. genome clusters can emerge from each other. Final trees for the mapped genome datasets were rooted manually according to the branching pattern in the 882 assembled *V. cholerae* core-genome tree, which diversity encompases that of the mapped genome trees.

From a subset of the 456 mapped 7PET genome alignments (n=335) corresponding to *Vc*H.9, a recombination-free phylogeny was inferred using ClonalFrameML v1.11^76^ with default parameters and using the ML mapped genome tree (restricted to the *Vc*H.9 genome tips) as a starting tree. BactDating^77^ v1.1 was then used to estimate a timed phylogeny (using 100,000 Monte-Carlo Markov chain iterations and otherwise default parameters) of the Yemen 2016-2019 genomes and relatives using the ClonalFrameML tree and day-resolved dates as input; median day of the year of isolation was used for isolates where these data were missing. Three independent chains were run from different random seeds and yielded close results.

Supporting data for phylogenetic analyses of the 882 assembled *V. cholerae*, 456 mapped 7PET genomes and 33 mapped *Vc*D genomes are avaible on Figshare repository at https://figshare.com/s/3fe31c131b00a2a08bb9 (doi: 10.6084/m9.figshare.16611823), https://figshare.com/s/4d83a32cce78a52b413e (doi: 10.6084/m9.figshare.16595999) and https://figshare.com/s/0be28064870c811120c5 (doi: 10.6084/m9.figshare.18304961), respectively.

### Correlation of spatio-temporal and phylogenetic distances

GPS data associated to the site of sample collection (health centres) were used to compute spatial geodetic distances using R script ‘gps_coords.r’^78,79^. Temporal distances were computed from the difference between day of collection (only available for 2018 and 2019 Yemen isolates). Phylogenetic distances were computed from the mapped genome tree using the function ‘cophenetic’ from the core R package ‘stats’^80^. Spatial, temporal and phylogenetic distances were compared using a Monte-Carlo approximation of the Mantel test as implement in the ‘mantel.randtest’ function from the R package ‘ade4’^81^, using 100,000 permutations to compute the simulated *p*-value. Maps showing the distribution of genomes clusters over the Yemen territory and in the region of Sana’a were obtained using QGIS 3.16.3 and the QuickOSM API to retrieve OpenStreetMap data, specifically level 4 administrative boundaries (governorates) in Yemen (last accessed 11 February 2021).

### Clade-specific SNPs and pangenome analysis

The synteny-aware pangenome pipeline Panaroo^82^ (v1.2.3) was run on the 882 assembled *V. cholerae* genome set with the option “--clean-mode strict” and default parameters otherwise. In parallel, a combined VCF file containing information on all SNP variation within the 456 mapped genome set was obtained using the ‘bcftools merge’ command. To identify clade-specific SNPs and accessory gene presence/absence patterns, we used custom R scripts^68^ to compare the combined VCF file and the gene presence/absence table output of Panaroo, respectively, to the mapped genome tree. Based on lists of genomes assigned to various clades and clusters (see Results), we identify SNPs or accessory genes that are specific of a focus clade in contrast to a background group or a sister clade, considering the contrast significant when the Bonferonni-corrected p-value is below 0.05 and when the frequency of an allele is above 0.8 in the focus clade and below 0.2 in the background clade, or conversely. Pangenome analysis files are available at https://figshare.com/s/675d2a9e424ad4f11646 (doi: 10.6084/m9.figshare.19519105). Putative anti-phage defense systems were searched by testing correlation of presence/absence patterns between ICP1-like phage and each pangenome gene cluster; only associations with Pearson correlation coefficients lower than -0.9 or greater than 0.9 and p-values lower than 10^−5^ were retained as significant.

## Data availability

Novel genomic data are available from the ENA/NCBI/DDBJ short read archive under the BioProject PRJEB34436. Four of the resulting assemblies comprised a single 123-kb contig corresponding to the ICP1-like phage; these assemblies were deemed uncontaminated and complete ICP1-like phage genomes and were deposited to GenBank under the accessions MW911612-MW911615. Complete hybrid genome assemblies for reference strains CNRVC019243 and CNRVC019247 were deposited to the ENA under the BioProject acessions PRJEB52123 and PRJEB47951 (Assemblies GCA_937000105 and GCA_937000115), respectively. Suplementary data are available online on the Figshare repository, under the following digital object ientifiers (doi): https://doi.org/10.6084/m9.figshare.16595999, https://doi.org/10.6084/m9.figshare.16611823, https://doi.org/10.6084/m9.figshare.18304961, https://doi.org/10.6084/m9.figshare.19097111, https://doi.org/10.6084/m9.figshare.19519105, https://doi.org/10.6084/m9.figshare.19575148.

## Acknowledgements

This research was funded in whole, or in part, by the Wellcome Trust (grants number 206194 and 108413/A/15/D). For the purpose of Open Access, the author has applied a CC-BY public copyright licence to any Author Accepted Manuscript version arising from this submission. This work was supported by *Institut Pasteur, Santé publique France*, and by the French Government’s *Investissement d’Avenir* program, *Laboratoire d’Excellence* “Integrative Biology of Emerging Infectious Diseases” (grant no. ANR-10-LABX-62-IBEID). We wish to thank Sally Kay, Joseph Woolfolk, Simon Clare and Charlotte Tolley for their support in sample management and logistics at WSI. We thank the WSI Pathogen Informatics team for support on informatics and the use of genomics anlaysis pipelines.

M.J.D. is an Official Fellow of Churchill College, Cambridge, and was previously supported by a Junior Research Fellowship at the College.

