## Supplementary Figures and Supplementary Text for "Genomic epidemiology of the cholera outbreak in Yemen reveals the spread of a multi-drug resistance plasmid between diverse lineages of *Vibrio cholerae*"

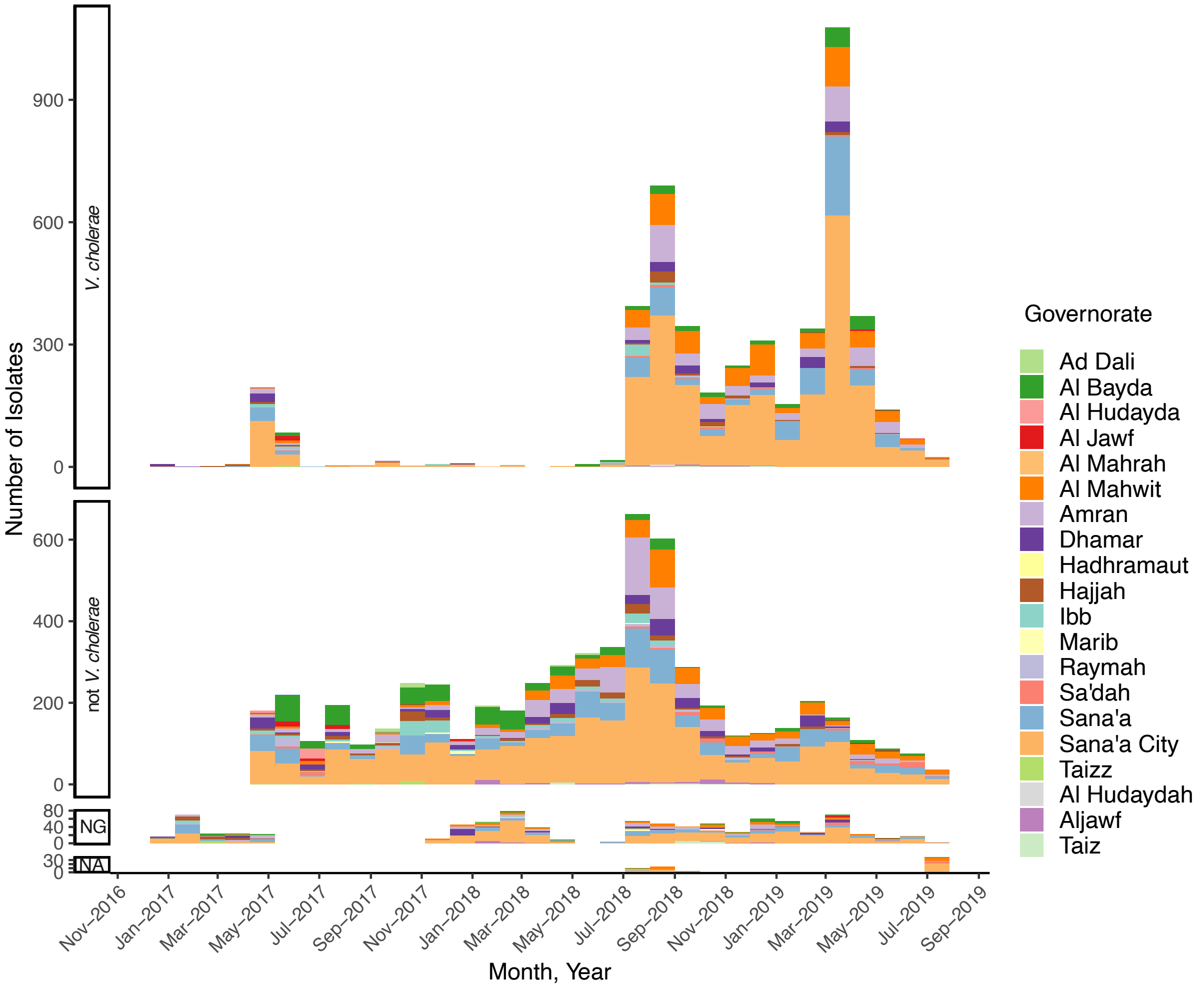


**Figure S1: Culture confirmation of samples derived from suspected cholera cases in Yemen, 2017-2019**

Distribution over time of *V. cholerae* culture result samples received at the NCPHL, broken down by governorate. Data are derived from Electronic Disease Early Warning System (eDEWS) linelists (Table S7). NG, no growth; NA, not available.

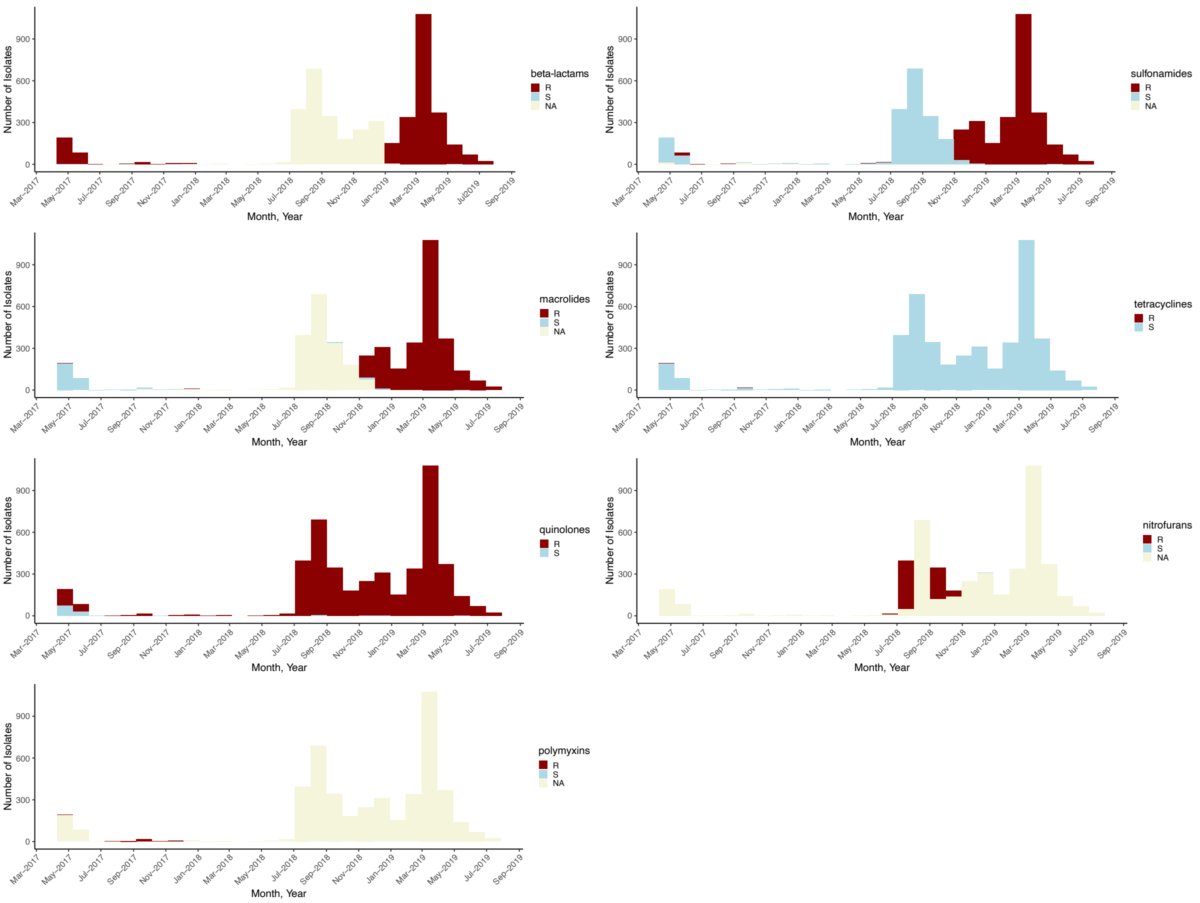


**Figure S2: Antibiotic susceptibility phenotypes of all *V. cholerae* isolates collected in Yemen, 2017 and 2019.**

Distribution over time of resistance and sensitivity to broad antibiotic classes among culture-confirmed *V. cholerae* isolates received at the NCPHL. Data are derived from eDEWS linelists (Table S7).

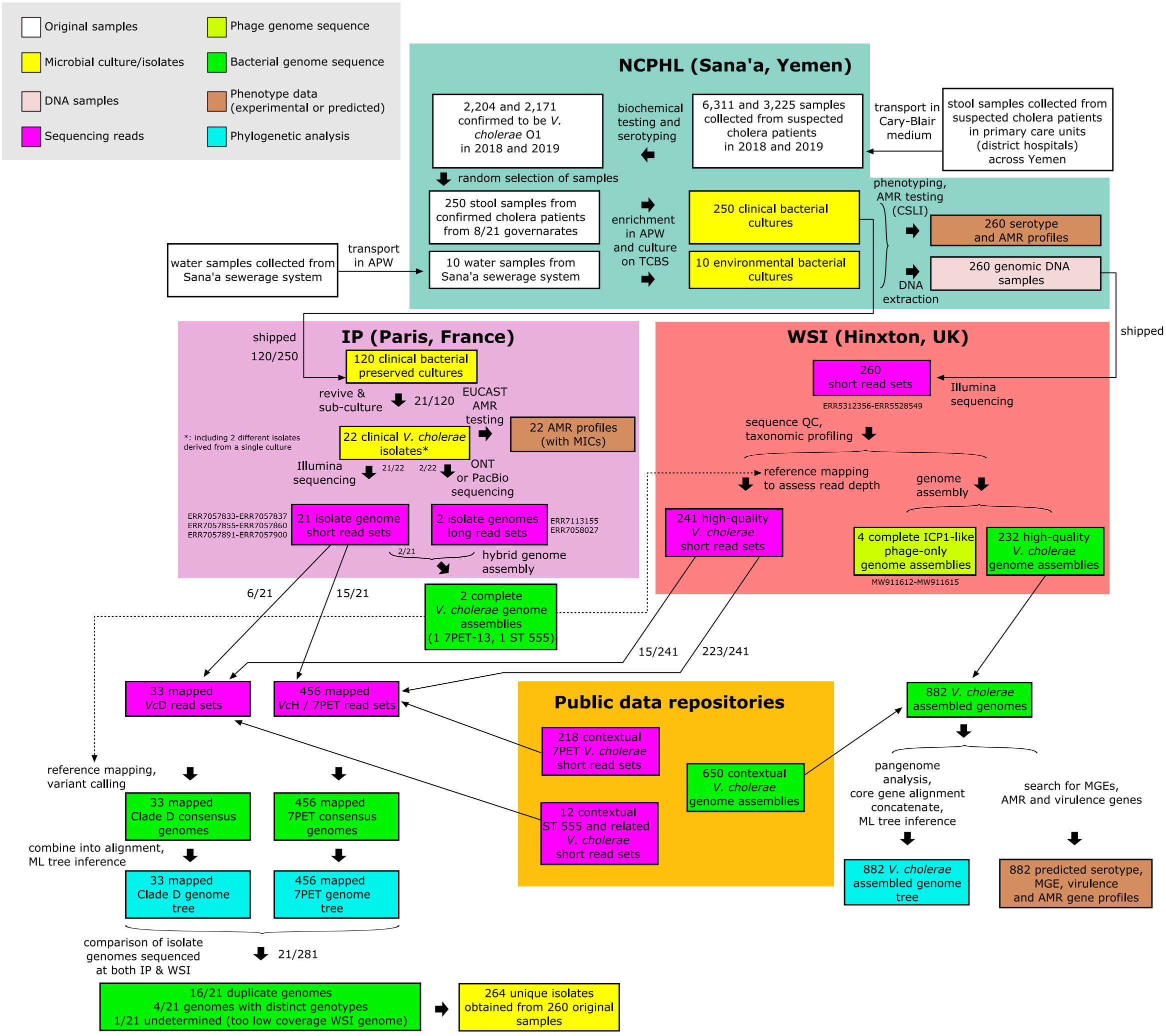


**Figure S3: Flowchart of sample collection, management and use in protocols and analyses** Experiments and analyses are itemized and grouped according to the different locations of the collaborative consortium were they were undertaken: NCPHL, The National Centre of Public Health Laboratories; IP, Institut Pasteur; WSI, Wellcome Sanger Institute.

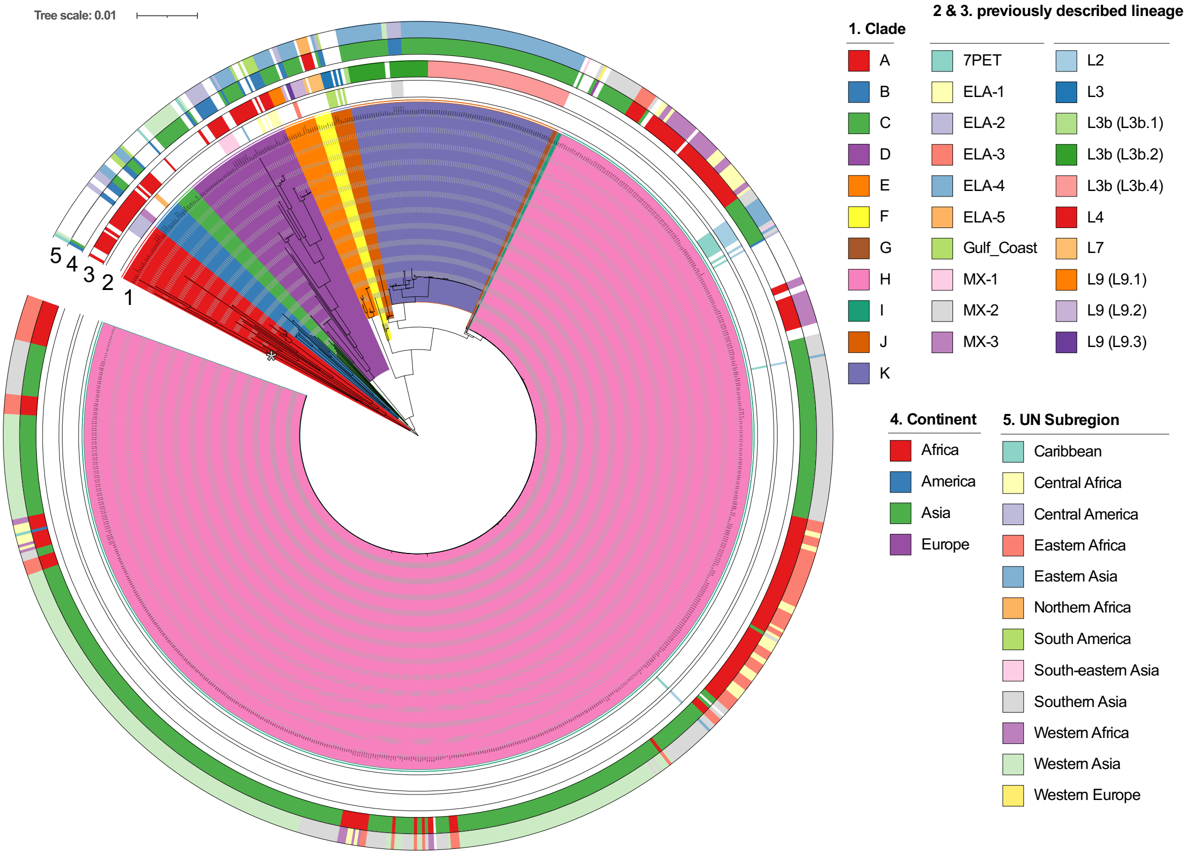


**Figure S4: Phylogenetic diversity of *Vibrio cholerae* isolates from Yemen and contextual samples**

Expanded version of phylogentic tree shown in Figure 1 (clades are not collapsed).

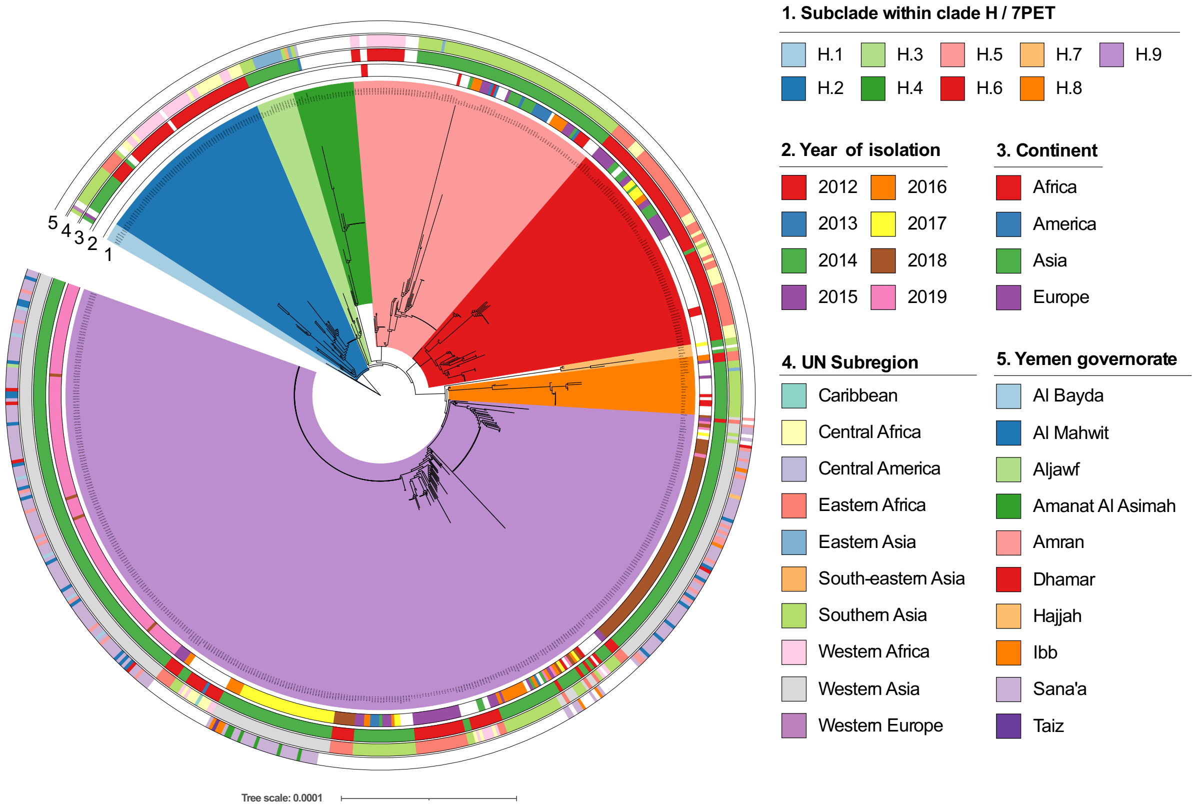


**Figure S5: Phylogenetic diversity of *Vibrio cholerae* *Vc*H isolates from Yemen and contextual samples**

Expanded version of phylogentic tree shown in Figure 1, focusing on the subtree of clade *Vc*H (collapsed in Figure 1), with details of the phylogenetic substructure.

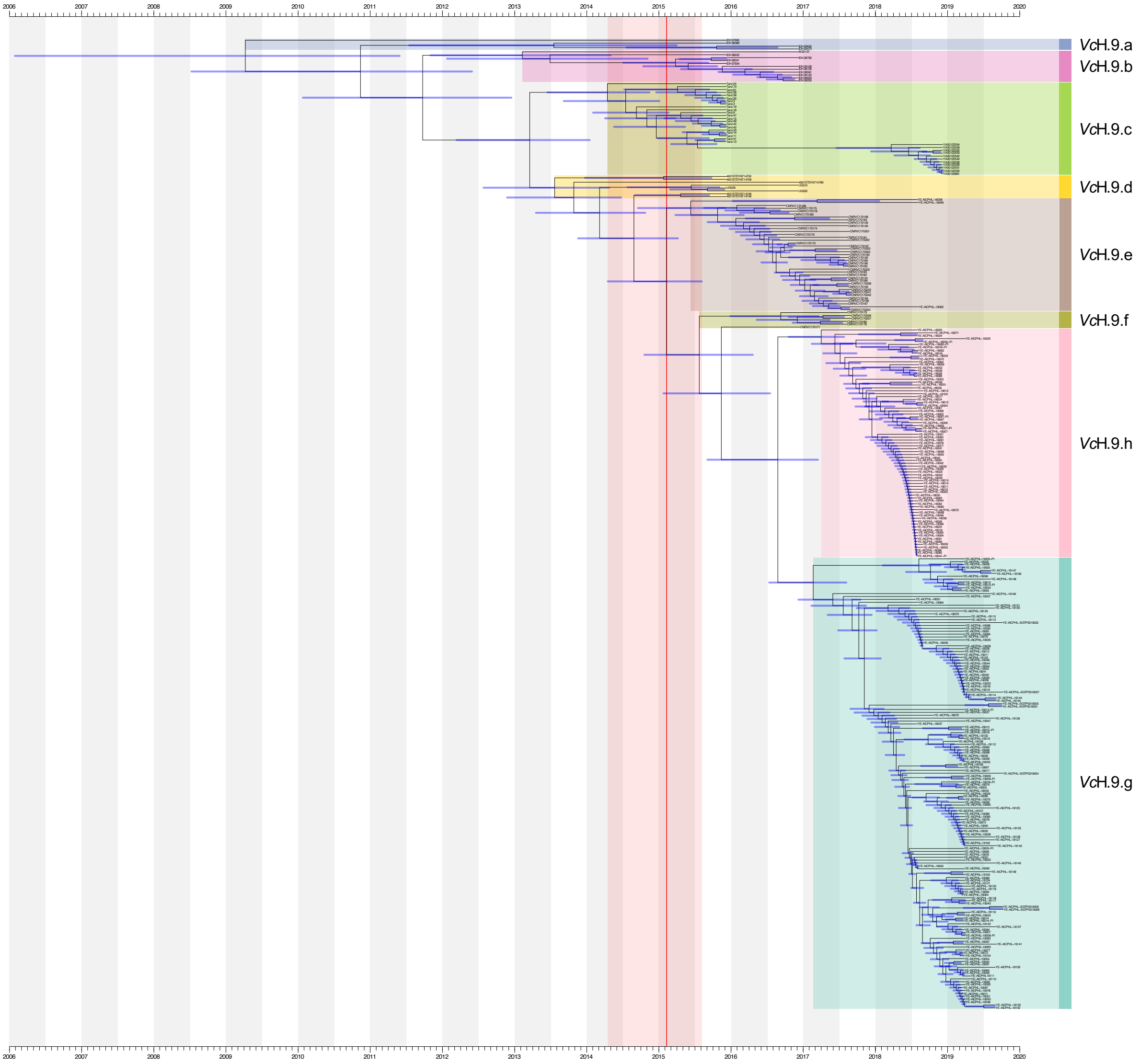


**Figure S6: Timed phylogeny of *Vibrio cholerae* *Vc*H.9 isolates from Yemen and contextual samples**

Timed phylogeny obtained by estimating the dates of nodes using BactDating[77] with the *Vc*H.9 subtree of the 456 mapped 7PET genome tree (as presented in Figure 2A) as input. Subclusters are labelled and coloured as per previous figures. X axis represents time in years.

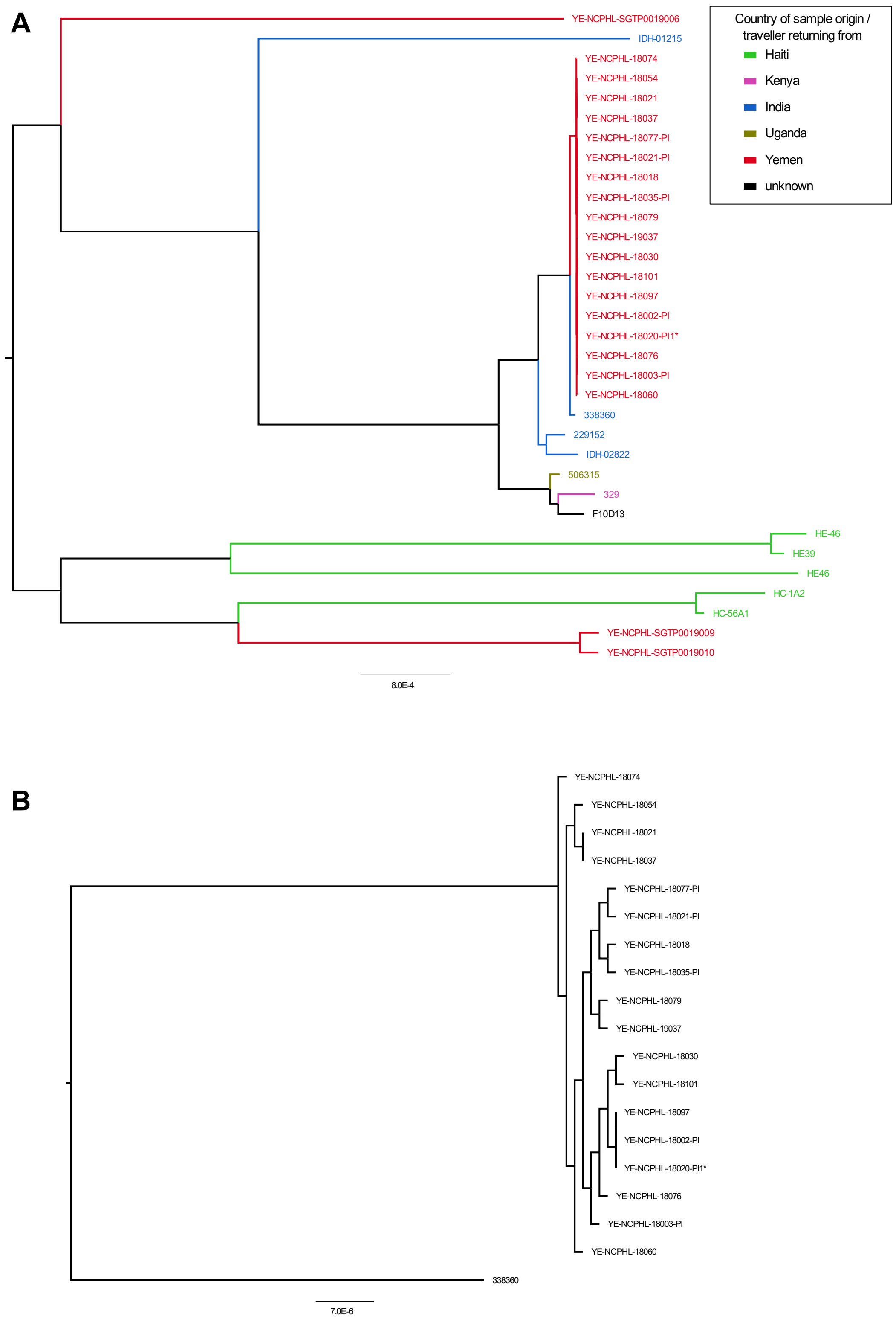


**Figure S7: Phylogenetic diversity of *Vibrio cholerae Vc*D/ST555 isolates from Yemen and contextual samples**

**A.** Maximum-likelihood phylogeny of the 33 mapped *Vc*D genomes. The tree was obtained based on the 91,312 SNP sites from concatenated whole-chromosome alignments. **B**. Subtree of the tree in A, focusing on the ST555 isolates from Yemen, and their closest relative, strain 338360.

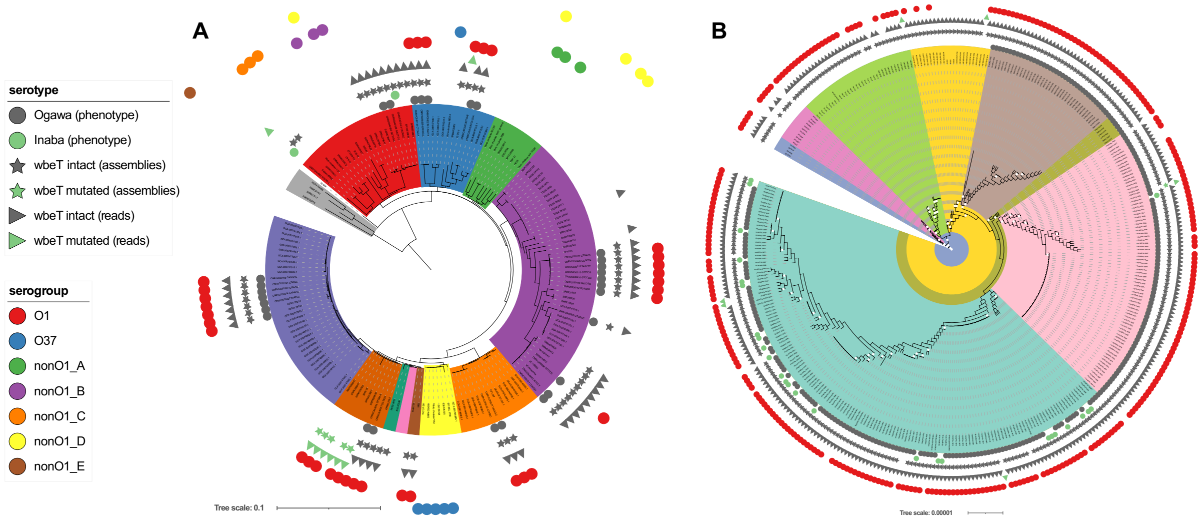


**Figure S8: *In silico* prediction of the O-antigen diversity among *Vibrio cholerae* from Yemen and contextual samples**

Prediction of LPS O-antigen serogroup and O1 serotypes projected on the isolate trees corresponding to **(A)** the core-genome tree as presented in Figure 1A and **(B)** the subtree of the mapped 7PET genome tree as presented in Figure 2A.

**Figure S9: Comparison of IncC plasmids pCNRVC190243 and pYA00120881**

Alignment of the differing regions of IncC plasmids pCNRVC190243 and pYA00120881 using Blastn. Regions of similarity are highlited by interleaving bands; band colour indicate similarity intensity and orientation of the alignment (blue, direct; pink, reverse). Figure was made using Easyfig[83].

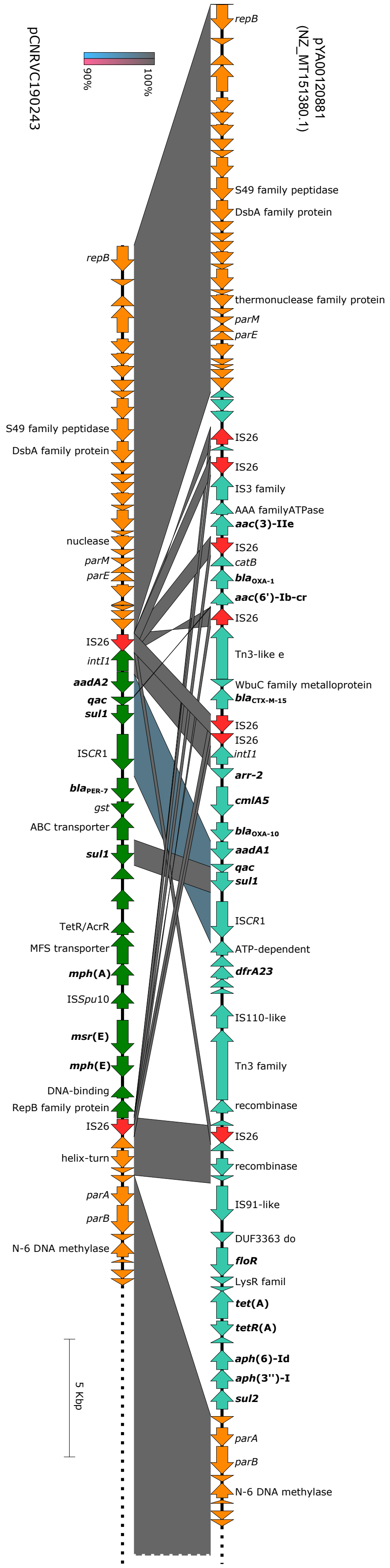

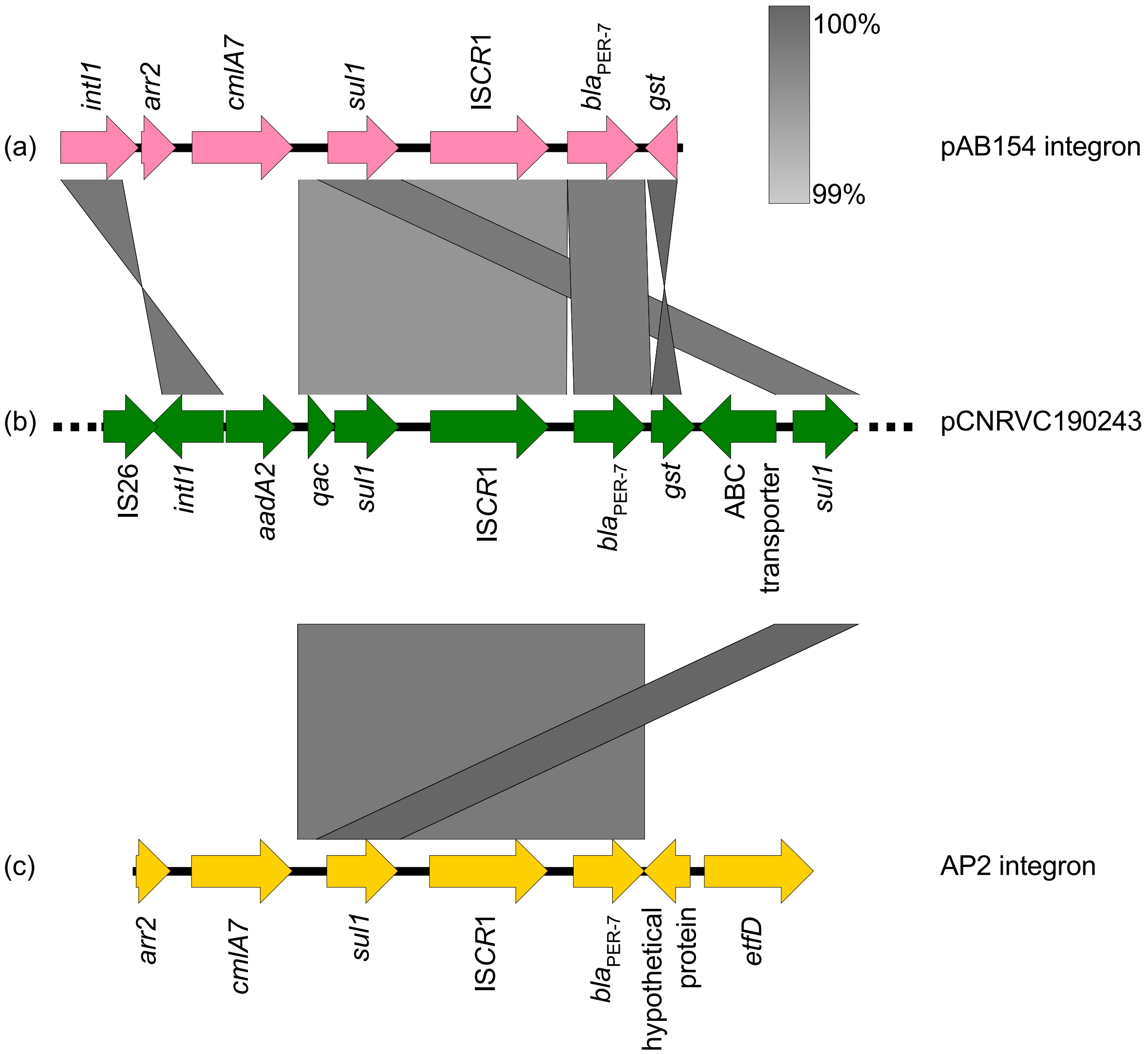


**Figure S10: Comparison of IS*CR*1 elements**

IS*CR*1 regions of (A) *A. baumanii* str. AB154 plasmid pAB154 integron (JQ639792.1), (B) pCNRVC190243/Yem*Vch*MDRI and (C) *A. baumanii* str. AP2 integron (HQ713678.1). Alignments were made using Blastn. Figure was made using Easyfig[83].

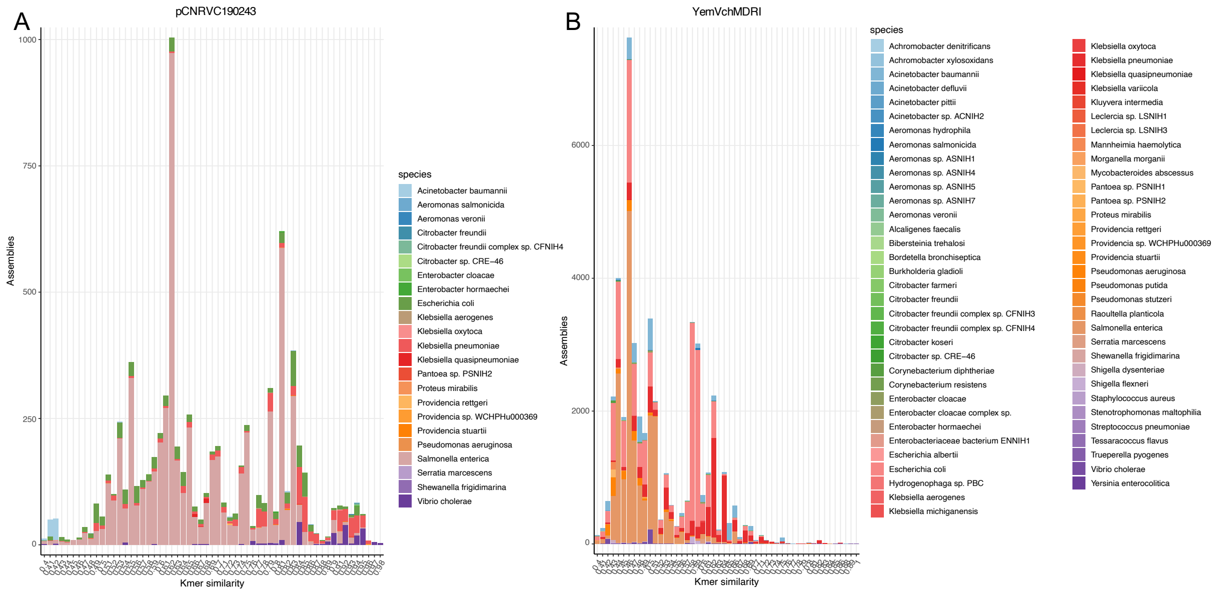


**Figure S11: Taxonomic distribution of sequences similar to pCNRVC190243 and Yem*Vch*MDRI elements**

Reference sequences of pCNRVC190243 and Yem*Vch*MDRI elements were searched using a COBS index against the 661k bacterial genome assembly database previously published by Blackwell *et al*.[58].

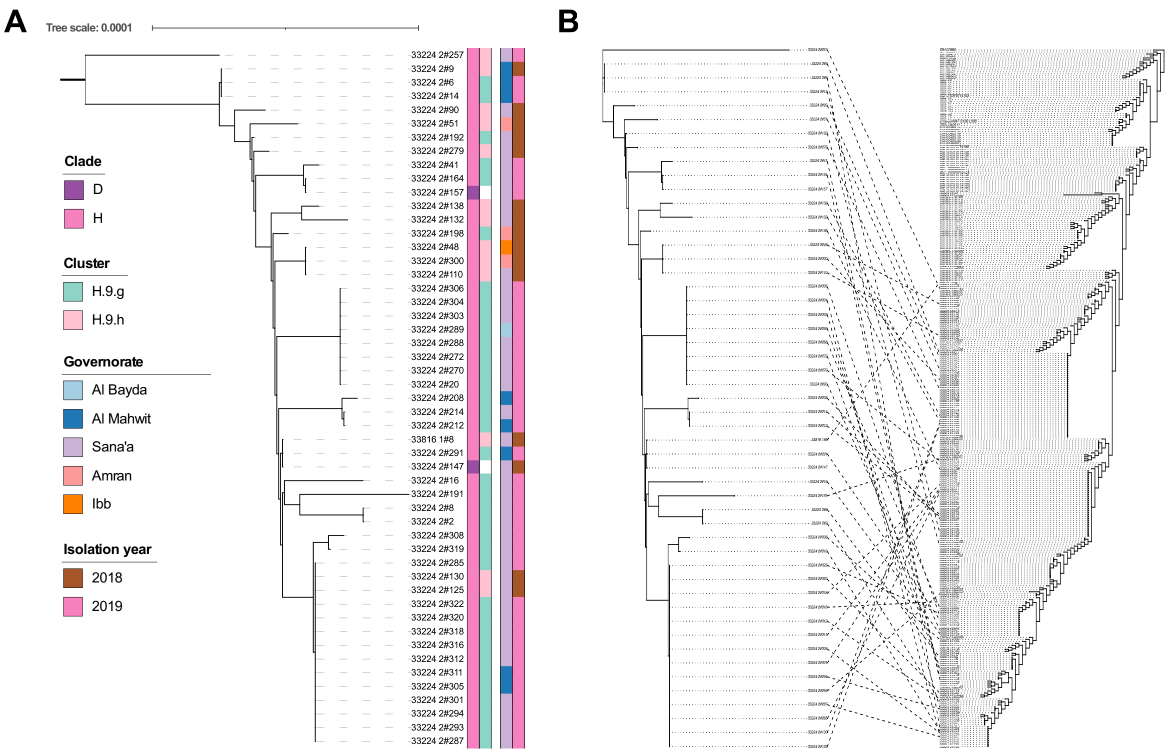


**Figure S12: Phylogenetic diversity of ICP1-family phage from yemen**

**A.** Phylogeny of ICP1-family phage sequences present in Yemeni *V. cholerae* sequencing read sets. These sequences were assembled by mapping reads to the most complete phage sequence obtained from the genome assembly of sample YE-NCPHL-19021 (GenBank accession MW911613.1). Clade and phylogenetic clusters (when part of *Vc*H.9) of the host *V. cholerae* genomes, as well as year and governorate of collection of the samples are indicated in couloured strips on the right (see key). **B**. Tanglegram facing the host bacterial phylogeny on the left (mapped genome tree as presented in Figure 2A) and the viral phylogeny on the right, both represented as cladograms. Dashed links match host and viruses originating from the same sample.

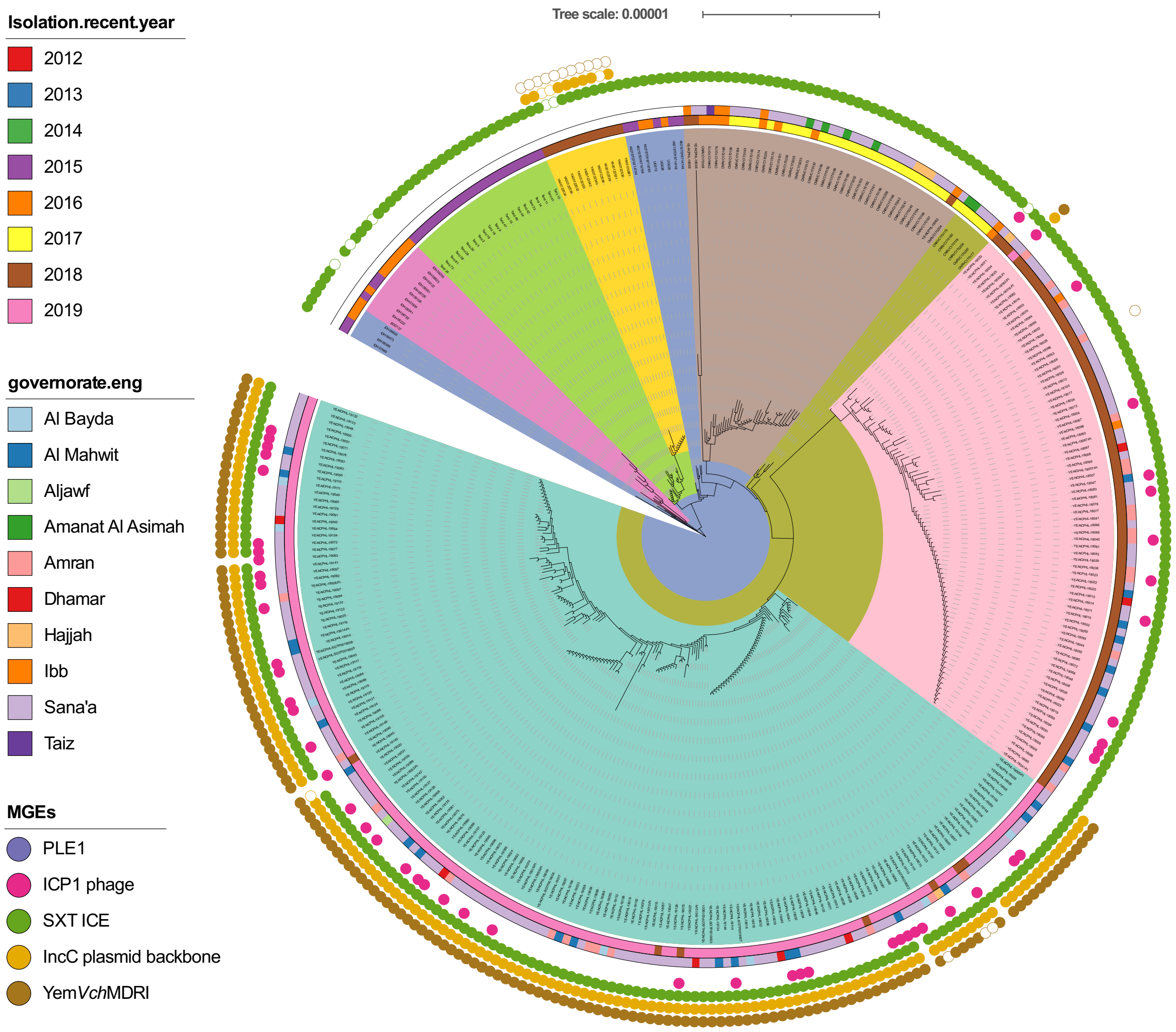


**Figure S13: Recombination-free phylogeny of *Vibrio cholerae Vc*H.9 isolates from Yemen and contextual samples**

Tree computed from the same alignment as in Figure 2A, but using ClonalFrameML[76] to infer the phylogeny reflecting the clonal propagation of the organism.

**Genomic epidemiology of the cholera epidemic in Yemen reveals the succession of epidemic clones driven by the acquisition of a multi-drug resistance plasmid**

### Supplementary Text

#### Samples and cultures

Out of the 4,375 samples confirmed to contain *V. cholerae* at the National Centre of Public Health Laboratories in the capital city Sana'a (NCHPL; Sana'a, Yemen), 260 cultures were used to prepare genomic DNA that were sent to the Wellcome Sanger Institute (WSI; Hinxton, UK). In parallel, an overlapping set of 120 live cultures were sent to the Institut Pasteur (IP; Paris, France), where only 21 could be revived (see Figure S3). Whilst samples received at both WSI and IP were derived from the same samples sharing the same nomenclature with the 'YE-NCHPL-' prefix, genomic and/or phenotypic information obtained from corresponding samples showed discrepancies (Table S13). Based on whole-genome phylogenies, we found that, out of 21 WSI/IP genome pairs, 10 had undistinguishable genotypes (branching as immediate relatives in the tree), and 6 branched separately on the tree but belonged to the same phylogenetic cluster – suggesting that the few observed genotypic differences could be ascribed to sequencing or mapping errors – confirming in 16 cases the genetic uniqueness of the twice-sequenced isolates. However, 4 genome pairs had completely distinct genotypes, belonging to cluster *Vc*H.9.h/7PET-T13 (all 4 genomes from strains isolated at NCHPL and sequenced at WSI) or *Vc*D/ST 555 (all 4 genomes from strains sub-cultured and sequenced at IP); the last genome sequenced at IP had no useable counterpart at WSI due to sequencing failures. This discrepancy for 4 samples may have stemmed from the presence of mixed strain cultures in the original samples resulting in different dominant clones being sequenced or subcultured in the respective recipient institutes. This hypothesis is supported by positive PCR screens for an O1 *V. cholerae*-specific marker (O1-type *rfb* operon)[1] from samples that resulted in non-O1/non-O139 strain cultures at IP but yielded O1 strain genomes at the WSI (data not shown). As a result, we chose to use two separate nomenclatures for samples received at WSI and IP, with the IP sample names bearing the '-PI' suffix (Table S2), whereas the WSI samples retained the initial name (Tables S1, S3, S4, S5; Figure 2A).

#### Spatiotemporal distribution of non-7PET *V. cholerae* isolates

The *V. paracholerae* and *Vc*D (ST1499) isolates obtained from Yemen had no close phylogenetic relatives taken from Yemen. When multiple closely related strains (ANI > 99.97%; Table S14) from these clades were observed, they were found in very localised areas: a pair of *Vc*K isolates (ST170) were isolated the same week of March 2019 from patients in Sana'a governorate, while a pair of *Vc*D isolates (ST1020) were taken on consecutive days in October 2019 from sewerage in Sana'a (Figure 1; Table S4). Finally, we characterised a cluster of 18 closely-related *Vc*D isolates belonging to ST555 (Table S6) – ten of which have high-quality genome assemblies (as presented in Figure 1 and Table S4) – which we found to differ from each otherby 0 to 10 SNPs (average 99.98% ANI similarity). Of these 18 isolates, 13 were isolated over a period of 11 days in late July/early August 2018, two at the end of August, two in October 2018, and one in March 2019 (Table S6). They were obtained from patients in the neighbouring governorates of Sana'a (n=7), Al Mahwit (n=4) and Amran (n=1), which are surrounding the capital city. Genomes from other ST555 isolates, including strains reported as linked to travelers returning from India in September 2015 and July 2016 (strains 229152 and 338360)[66], as well as closest relatives from our core-genome tree, were gathered to build a tree of mapped *Vc*D genomes using the 2018 Yemen strain CNRVC190247 complete genome as a reference (Figure S7). The closest relative to Yemeni ST555 isolates, isolate 338360, differs from the *Vc*D ST555 genomes sequenced here by between 763-800 SNPs, ruling out direct clonal relationships.

**Inconsistencies between phenotypic characterisation and *in silico* predictions of O-antigen serogroup, serotype and AMR profile**

Phenotypic characterisation of *V. cholerae* isolates at NCHPL included serotyping, testing with antisera for O1 serogroup serotypes Inaba and Ogawa. The isolates corresponding to the 260 novel Illumina genomes reported in this study were all positive for either Ogawa (n=230) or Inaba (n=30) (Table S1). However, these results proved inconsistent with our in-silico serotyping using ARIBA (see Methods), showing that only 218/241 testable genomes were of serogroup O1, and among them only two were predicted to be of the Inaba serotype (isolates YE-NCPHL-18053 and YE-NCPHL-19014 = YE-NCPHL-19014-PI = CNRVC190243) (Table S5). Serotyping conducted on the subset of the samples sub-cultured at the IP (Figure S3) showed a higher degree of consistency with the *in silico* serotype predictions, with only strain YE-NCPHL-19015-PI serotyped as Inaba while *in silico* serotyping of corresponding genome assembly for isolate YE-NCPHL-19015 predicted an Ogawa serotype (Table S2).
In addition, antibiotic sensitivity testing at NCHPL showed an ampicillin resistant phenotype for all tested strains, including those not carrying the pCNRVC190243 plasmid or without any other potential resistance determinant as predicted using Abricate (see Methods; Table S2); and ampicillin sensitivity was confirmed for all seven 2018 (plasmid-free) 7PET-T13 genomes cultured at IP (Table S2). Similarly, some 2019 isolates were found resistant to doxycycline (n=11) or tetracycline (n=2), or showed resistance to ciprofloxacin (n=17), but none of these phenotypes could be supported by genotype predictions with Abricate, nor were they confirmed when cultured at the IP (YE-NCPHL-19005-PI and YE-NCPHL-19008-PI are $TET^S$, YE-NCPHL-19009-PI is $CIP^S$; Table S2).
We thus consider that it is possible that reagent expiry at NCHPL may have contributed to the discrepancies between phenotypic and genotypic serotyping as well as ampicillin, tetracycline and ciprofloxacin resistance prediction.

On the other hand, the Abricate screen found that all 7PET-T13 Yemen isolates carried the *tet*(34) and *catB9* genes (Table S4) and were thus predicted to be resistant to tetracycline and chloramphenicol. However, *catB9* gene is known not to provide chloramphenicol resistance in this genomic background (Weill et al., Science 2017), consistent with chloramphenicol susceptibility of strains tested at IP (Table S2). Neither tetracycline resistance nor oxytetracycline resistance was observed in phenotypic tests conducted at NCHPL and IP

(Tables S1, S2), even though *tet*(34) has been described to only provide oxytetracycline resistance in a $Mg^{2+}$-rich medium[2], a condition that may not be met during the phenotypic tests. However, the gene *tet*(34) was described to be similar to the xanthine–guanine phosphoribosyl transferase gene of *V. cholerae*[3], and was detected in 881/882 of our assembled *V. cholerae* genome collection (Table S4), suggesting that it is a core gene of the species, and that any encoded tetracycline resistance phenotype would be shared across the species – which has never been observed; we therefore conclude that this *tet*(34) gene is a false positive hit not related to tetracycline or oxytetracycline resistance.
Finally, aminoglycoside resistance is predicted for a range of strains with diverse genotypes, based on the presence of genes *aadA2*, *aph*(3'')-Ib and *aph*(6)-Id. However, no resistance to antibiotics of this class was observed in tests conducted at both NCHPL and IP, with all strains tested for sensitivity to kanamycin, amikacin, gentamycin reported susceptible. The reported genes, including *aadA2* located on the YemVchMDRI PCT, may not be functional, or may not be expressed in the tested conditions.

Given that serotype and antibiotic susceptibility test results obtained on the lower-size sample at the IP generally agreed with our genotype-based *in silico* predictions, we assumed NCHPL test results may have suffered from adverse conditions leading to unreliable results, and we therefore chose to report genotype-based predictions for these traits, except for false positive predictions of resistance to tetracycline, chloramphenicol and aminoglycosides.

**Genomic context of the standalone MDR pseudo-compound transposon**
In the ten assembled genomes of *V. cholerae* ST555 (VcD) isolates, the MDR pseudo-compound transposon (PCT) YemVchMDRI is present without the IncC plasmid backbone observed in other genomic backgrounds of 2018-2019 Yemeni *V. cholerae* isolates. In the ST555 background, represented by the complete genome sequence of strain CNRVC190247 (= YE-NCPHL-18035-PI), the PCT is inserted in chromosome 2, between:
- (in 5' of the PCT, referring to the orientation on pCNRVC190243) two transposases (CNRVC190247_02871 and CNRVC190247_02872), followed by a large ORF (CNRVC190247_02873) encoding a 1273-amino acids hypothetical protein, that is conserved among ST 555 genomes; the following segment begins with a gene encoding a protein of the NirD family (CNRVC190247_02875), which is part of the *V. cholerae* species core-genome.
- (in 3' of the PCT, referring to the orientation on pCNRVC190243) a gene encoding a cobyric acid synthase (CNRVC190247_03766) and an operon coding for a molybdenum ABC transporter (CNRVC190247_03763-03765); this region and the following segment are also part of the *V. cholerae* core genome.

In all Illumina short-read assemblies of ST555 genomes, YemVchMDRI appears as a standalone contig. Taking the genome assembly of strain YE-NCPHL-18060 (assembly id 33224_2#171) as an example, the entire contig #32 (long of 20,249 bp) corresponds to the PCT, except for the IS26 elements that should be located at both ends of the PCT (forming direct repeats). A scaffold assembly graph (based on the paired-end read information) however reveals that the PCT/contig #32 is connected at both ends to a contig which represents the IS26 element (contig #54), thus confirming the expected structure of the PCT.

The IS26/contig #54 is itself connected to multiple contigs: contig #14 (106,332 bp) at one end, which corresponds to the region located upstream the PCT on chromosome 2 of the reference strain CNRVC190247 genome; at the other end, through ramifications of the assembly graph, contig #54 is connected to contigs #11, #24, #29, #31, #36 and #45, none corresponding to regions located next to the PCT in reference strain CNRVC190247 genome, indicating that the short read-based assemblies cannot resolve the structure of this repeat-rich region. This pattern is replicated across all short-read assemblies of Yemeni ST555 strains. Despite uncertainty on what locus lies downstream of the PCT in short read-based assemblies of ST555 strains, the identity of the upstream regions with that of the hybrid assembly of reference strain CNRVC190247 provides reasonable support for the PCT being inserted at the same chromosomal locus in all Yemeni ST555 strains.

**Phylogenetic hypothesis testing**

From the ML tree (output of RAxML-NG) computed for the SNP alignment of 457 mapped 7PET genomes, some relationships between genome clusters appeared to be well-resolved (very low branch bootstrap supports) and indicated some alternative topologies would be equally supported. In particular, we considered the distribution of genomes in the initial ML tree to be not parsimonious with regard to the associated metadata. Indeed, a cluster gathering all genomes of isolates sampled from Kenya in 2010-2015 and Uganda in 2014-2016 (cluster *Vc*H.9.d in the final tree) was paraphyletic, due to the inclusion of a cluster of genomes of isolates sampled from Tanzania in 2015 and Zimbabwe in 2018 (cluster *Vc*H.9.c in the final tree). Similarly, the cluster containing most of the Yemeni isolate genomes from 2018 Yemeni isolates (cluster *Vc*H.9.h in the final tree) emerged from the clade containing most of the Yemeni isolate genomes from 2019 Yemeni isolates (cluster *Vc*H.9.g in the final tree). In both cases, the included clade was connected to the root of the inclusive clade by a string of branches with very low bootstrap supports. We therefore formulated hypotheses where the included clades (clusters *Vc*H.9.d and *Vc*H.9.h, respectively) would branch as sister clades of the initially inclusive clades (clusters *Vc*H.9.c and *Vc*H.9.g, respectively). We first tested the relationship between *Vc*H.9.h and *Vc*H.9.g clusters, and obtained nearly equal support for the inclusion of *Vc*H.9.h within VcH.9.g clade (log likelihood = -5719994.708326), or their branching as sister clades (log likelihood = -5719994.708058); the latter hypothesis received slightly higher support (Shimodaira-Hasegawa [SH] test *p*-values: 0.4629 and 0.5125, respectively) and was used as background for testing the relationship between *Vc*H.9.c and *Vc*VcH.9.d. We obtained significantly stronger support for the branching as sister clades of *Vc*H.9.c and *Vc*H.9.d (log likelihood = -5719994.701928) when compared to the initial inclusion hypothesis (SH test *p*-values: 0.4871 and 0.0001, respectively). When the three hypotheses are compared together, the support is overwhelmingly in favour of the tree with both cluster pairs resolved as sisters (SH test *p*-values: 0.0001 for both hypotheses with *Vc*H.9.d included in *Vc*H.9.c, and 0.5167 for the sister-clade hypothesis); this latter tree topology is described in the main text and represented in Figure 2.

**ICP1-like lytic bacteriophage**

We detected the presence of an ICP1-family of lytic bacteriophage (ICP1) in 51/232 Yemeni *V. cholerae* genome assemblies, where they accounted for > 1% of sequencing reads per sample (Tables S3, S4). Occurrence of phage genomes in the sequenced genomic DNA from bacterial isolates likely reflects contamination of the isolate cultures with phages present in the original stool or sewage samples (see Methods). Some read sets showed high levels of contamination with phage-derived reads (16 samples with >50% ICP1 reads), suggesting that ICP1 phages may have infected the bacterial isolates in the samples and undergone several lytic cycles. The 51 assembled phage sequences showed in average 5.94 pairwise SNP differences (nucleotide diversity $\pi$ of $4.87 \cdot 10^{-5}$), a variation two orders of magnitude lower than seen previously within a collection of diverse ICP1-family phages isolated over a 12-year period in Bangladesh[39]. The closest matches to the Yemeni ICP1 were found in the genome data for *V. cholerae* strain THSTI_45869, isolated in Delhi, India in 2015[11] (94% coverage and 99.7% nucleotide identity), and an ICP1 phage (MN402506.2) isolated in Chandigarh, India in 2015 from community sewage water (94% coverage and 99.9% identity; Chaudhary, N. and Taneja, N., direct submission to GenBank).

Phages were mostly found associated with *V. cholerae* O1-serogroup *Vc*H.9 strains (Figure S12A), consistent with the previously reported receptor specificity of this phage family[74], but were also associated with two non-O1/non-O139 clade D strains (Tables S6). Presence of ICP1-like phage sequences in these genomic read sets (6.5% and 5% reads, respectively) suggests the page was present and replicating in these samples, but it is possible that a bacterial strain mixture including an O1-serogroup strain occurred in the original samples, as was observed for other samples (see Supplementary Text). ICP1-positive samples originated from five governorates, mostly from Al Mahwit and Sana'a (Tables S1,S3). The ICP1-like phage phylogeny showed limited geographic and temporal clustering, potentially reflecting a pattern of local transmission (Figure S12A). In addition, the viral phylogeny showed a branching structure incongruent with that of the bacterial host (Figure S12B, focusing on the *Vc*H.9 subtree), suggesting multiple horizontal transfers of the phage between host bacteria, consistent with the known lytic mode of transmission of ICP1-like phages.

Looking into genetic determinants of phage sensitivity that could explain variable phage occurrence in samples, we tested the significance of correlation of occurrence of the phage against presence of a range of known anti-phage defence systems or, without *a priori* targets, against all other pangenome genes, but found no significant association with any gene (see below). Instead the variable occurrence of the phage may result from the stochasticity of culture contamination during the process of bacterial isolation from the original samples. The recovery of an ICP-1-like phage in association with almost a quarter of *V. cholerae* isolates in Yemen in 2018-2019 suggests that this virus could often accompany the 7PET populations, as was previously observed in South Asia[39], and reinforces the view that these phages may represent biomarkers for 7PET *V. cholerae* populations[75].

**Testing association of ICP1-like phages with other genomic traits**

To investigate whether the variable presence of the ICP1 among Yemeni 7PET-T13 isolates could be explained by increased sensitivity or resistance conferred by some genetic determinants, we tested the significance of correlation of presence of the phage against occurrence of a range of other traits.

First, we looked into the association with the presence or absence of known anti-phage defense systems, none of which were significant: all *Vc*H genomes featured an intact ICEVchInd5 Hotspot 5 and none of the Yemeni host genomes carried the PICI-like elements (PLE) – both elements known to provide protection to the bacterial cell against bacteriophage infection or dissemination[5,6]. In addition, CRISPR-Cas loci were found in only four environmental Yemeni isolates, carrying one or two loci of type I-F[7], all located within or next to a prophage locus (Table S4). Three of these CRISPR-Cas-positive isolates belonged to *Vc*D (but outside ST 555; Table S6), one of which was ICP1-positive, and one (YE-NCPHL-SGTP0019002) belonged to 7PET-T13 cluster *Vc*H.9.g, and was not ICP1-positive (Table S5). The latter occurrence could represent a novel finding as the presence of a CRISPR-Cas system has never been reported previously for 7PET isolates; in this 7PET isolate genome, the contigs carrying the CRISPR-Cas systems (contigs #28 and #29) both have landmarks of mobile elements such as integrases, suggesting they constitute a large Tn*7* transposon and a prophage, respectively, and show 100% homology with segments of larger contigs from genomes of *Vc*D isolates YE-NCPHL-SGTP0019009, suggesting these mobile elements may have been recently transferred from a *Vc*D background to a 7PET background. However, we cannot rule out that this occurrence of CRISPR-Cas-carrying MGEs in this 7PET isolate genome could stem from contamination by another genome, typically that of an endemic *Vc*D strain, or by the free form of the MGEs; occurrence of the same CRISPR-Cas-carrying elements in several other environmental sewage samples suggests such contamination could easily happen. Of note, the CRISPR-Cas-containing contigs of YE-NCPHL-SGTP0019002 genome are both only showing a single-end connection in the assembly graph (i.e. one end has no link to any other part of the genome based on paired-end read information), while the connected end links to a short repeated sequence; this again suggests these contigs were derived from standalone segments of DNA that did not belong to the strain's genome.

We further looked for other unknown anti-phage defence mechanism using correlation of pangenome genes against ICP1-like phage presence, but no gene – except the ICP1 phage's own genes – showed any strong pattern of (anti-)association with phage presence (data not shown).
